# *Undaria pinnatifida* exudates trigger shifts in seawater chemistry and microbial communities from Atlantic Patagonian coasts

**DOI:** 10.1101/2020.10.21.349233

**Authors:** Mariana Lozada, María C. Diéguez, Patricia E. García, Gregorio Bigatti, Juan Pablo Livore, Erica Giarratano, Mónica N. Gil, Hebe M. Dionisi

**Author notes:** Corresponding author: Mariana Lozada, IBIOMAR-CONICET, CCT CONICET-CENPAT, Bvd. Brown 2915, U9120ACD Puerto Madryn, Chubut, Argentina.

## Abstract

The invasive kelp *Undaria pinnatifida* has spread from northeastern Asia to temperate coastal environments worldwide, with profound effects on colonized ecosystems. In this work, we analyzed the effect of exudates from *U. pinnatifida* on the chemical and microbial properties of seawater from a semi-enclosed gulf from Atlantic Patagonia. Exudates of *U. pinnatifida,* consisting mainly of carbohydrates, were released at a rate of 1.6 ± 0.8 mg C g^−1^ algae day^−1^, affecting the quality and optical properties of seawater in experimental incubations. Parallel factor analysis based on excitation-emission matrices collected from exudates revealed the presence of two humic-like and one non-humic fluorescent components. Exudate release stimulated microbial growth and polysaccharide degrading activity in seawater. After a 7-day incubation of fresh seawater with the exudates, changes in microbial community structure were analyzed by large-scale 16S rRNA gene amplicon sequencing. Copiotrophic and fermentative genera such as *Spirochaeta* (Spirochaetes) and *Propionigenium* (Fusobacteria) increased in the incubations with algal exudates. Genomic potential prediction revealed that the selected bacterial community could have higher ribosome content - an indicator of the potential for reaching higher metabolic rates - and genes for the degradation of complex organic compounds such as polysaccharides and other carbohydrates present in the exudates. Nutrient addition triggered the emergence of other microbial populations with different ecophysiological niches: unclassified Flavobacteriales, unclassified bacteria related to the recently described Phylum Kiritimatiellaeota, as well as potential pathogens such as *Vibrio* (Gammaproteobacteria) and *Arcobacter* (Epsilonproteobacteria), suggesting potential synergistic effects between invasive macroalgae and human activities.

## Introduction

The Asian kelp *Undaria pinnatifida* (Phaeophyceae, Laminariales) is an invasive seaweed that has aggressively spread over the last decades through Northern Europe, the Pacific coast of North America, the coast of Australia and New Zealand, and the South Atlantic coast (Minchin and Nunn 2014; South et al. 2017). On the South Atlantic Patagonian coast (Argentina), it was first recorded in 1992 on a pier of Puerto Madryn city (42.75°S), located in the Nuevo Gulf. Since then, it has expanded both southwards and northwards, covering almost 10° of latitude along the argentinean coast (Bunicontro et al. 2019). *U. pinnatifida* forms dense seasonal forests in coastal waters up to 15 m depth, and it is considered an autogenic ecosystem engineer due to its capacity of significantly altering its surrounding environment (Casas et al. 2004; Raffo et al. 2009; Irigoyen et al. 2011b). In the semi-enclosed waters of the Nuevo Gulf, this species has already displaced other algal genera like *Codium* and *Dictyota* (Raffo et al. 2009), and affected sites used for commercial and recreational activities such as aquaculture, fishing and diving (Irigoyen et al. 2011a). Native consumers such as sea urchins do not seem to be able to control its recruitment (Teso et al. 2009). In turn, activities impacting coastal rocky reef communities such as scuba diving may further enhance the settlement of this species, setting a detrimental feedback loop (Bravo et al. 2015).

Living brown algae exudate important amounts of polysaccharides, proteins, lipids and aromatic compounds that collectively contribute to seawater dissolved organic matter (DOM) pool, increasing dissolved organic carbon (DOC) concentration and shifting its quality (Sieburth 1969; Abdullah and Fredriksen 2004; Wada and Hama 2013). These variables are in turn key drivers of several physical and chemical water properties (light penetration, temperature, pH, nutrients) that shape the structure and functioning of coastal food webs. For example, light absorption by the chromophoric fraction of DOM (CDOM) controls light penetration in the water column, determining underwater visibility and regulating the primary production of phytoplankton and other algal species (Blough, N. V. and Del Vecchio, R. 2002; Mostofa et al. 2013). In addition, DOM along with nutrients exert a bottom-up control on the microbial loop and, depending on the amount and chemical nature of the released substances, may have a significant impact on the planktonic food web (Kujawinski 2011). In turn, the humic fraction of DOM affects the fate, bioavailability, and toxicity of pollutants in aquatic systems (Artifon et al. 2019).

Polysaccharides are part of the labile fraction of algal exudates, thus they constitute a readily available substrate for bacteria, fueling both pelagic and attached microbial communities (Abdullah and Fredriksen 2004; Bengtsson et al. 2011; Haas et al. 2011). Algal polysaccharides vary greatly in their monosaccharide constituents and glycosidic linkages (Deniaud-Bouët et al. 2014), and in turn, this chemical diversity selects for a variety of bacterial populations capable of assimilating specific polysaccharides through the expression of enzymes, transport systems and transcriptional regulators (Arnosti 2014; Grondin et al. 2017). Two polysaccharides, alginates and fucoidans, comprise the major fraction of the matrix that forms brown algae exudates (Kloareg and Quatrano 1988). The first step of the assimilation process is the depolymerization of these polysaccharides by the extracellular enzymes alginate lyases (Matos et al. 2016) and fucoidanases (Kusaykin et al. 2016).

Despite the central role of macroalgal-derived DOM in coastal biogeochemical processes through its impact on local microbial communities, little is known about the composition of seaweed exudates, their contribution to the total C budget and their role in shaping seawater microbial communities. The effects of algal DOM on bacterial communities of coral reefs have been studied in detail, unveiling the mechanisms by which reef health is affected (Smith et al. 2006; Nelson et al. 2013). This led to the hypothesis of “microbialization”, a process by which labile DOM from algae associated to nutrient pollution stimulates microbial communities, unbalancing the whole coral reef system (Haas et al. 2016). Moreover, sugars similar to the ones present in exudates have been shown to promote the expression of virulence factors and other interaction proteins in microbial communities of coral reefs, with negative consequences on water quality (Cárdenas et al. 2018). While the effects of *U. pinnatifida* on the upper trophic levels of the invaded communities have been thoroughly studied (Teso et al. 2009; Raffo et al. 2009; Irigoyen et al. 2011b), no information is yet available regarding the potential effects of this invasive species on seawater physicochemical properties or microbial community structure. In this investigation, we hypothesized that exudates of *U. pinnatifida* could change seawater properties, with consequences for the associated microbial communities. Compared to other algae, the larger size and higher biomass of *U. pinnatifida* may have an effect on the carbon dynamics of invaded areas. DOM stimulation of the microbial loop, a key component of the planktonic food web, may propagate to higher levels of the food web potentially affecting the biogeochemical properties of the entire ecosystem.

The aim of this work was to empirically analyze the effects of *U. pinnatifida* exudates on seawater and its associated bacterial communities in a Patagonian rocky shore environment (Punta Este, Chubut, Argentina) extensively impacted by this invasive species. In our experimental approach, *U. pinnatifida* were first incubated in controlled conditions of temperature and photoperiod to obtain and analyze the exudates. As a second step, algae were removed and the exudates were mixed with fresh seawater to further study changes in DOM and in microbial community structure.

## Materials and Methods

### Sampling

Algae and seawater samples were obtained from Punta Este (PE, Chubut, Argentina, 42°47′ S; 64°57′ W). PE is a shallow hard bottom shore located ~5 km south of Puerto Madryn city, in Nuevo Gulf. Previous studies in this site have shown high biomass of *U. pinnatifida* (ca. 443 g m^−2^), especially at low depths (ca. 5m) (Rechimont et al. 2013), a higher biomass than the one yielded by algae such as *Codium vermilara* (ca. 82 g m^−2^) and *Dictyota dichotoma* (ca. 23 g m^−2^) inhabiting the same zone (Rechimont et al. 2013). Sporophytes of approximately 50 cm length and water samples were obtained from 5 m depth on the rocky shores of PE, between August and September 2019, in the period of highest biomass of the *U. pinnatifida* cycle. Seaweed and water samples were collected fresh and transported immediately to the laboratory.

### Experimental setup

*U. pinnatifida* individuals were allowed to exudate under controlled conditions (experiment 1), after which the algae were removed and mixed with fresh seawater obtained from PE, with and without nutrient addition (experiment 2).

For experiment 1, blades were cut from sporophytes, and placed for 3 days for healing and acclimation in tanks under controlled conditions (14°C, 33.5 ‰ salinity and 12:12 h light:darkness photoperiod). Blades (hereinafter referred to as “algae”) were placed in closed transparent plastic containers filled with exactly 2 L of serially filtered (100, 10 and 1 μm) and UV-sterilized seawater, with a relation of approximately 5% algal biomass/seawater (algae wet weight range: 95-113 g). Algae were further incubated for 48 h in the same conditions provided during the acclimation. Three replicates were setup with algae (EX) and three with filtered and UV-sterilized seawater without algae (incubation controls, INC). At each sampling time (0, 24, and 48 h), 200-ml liquid samples were obtained from the containers for subsequent chemical and microbiological analyses.

After 48 h, algae were removed, and exudates from experiment 1 were mixed with fresh seawater from PE site to set up a second experimental incubation (experiment 2). The design of experiment 2 was based on the work of Haas and collaborators (Haas et al., 2011), which was in turn based on the “seawater dilution culture incubation technique” as developed by Carlson et al. (Carlson et al., 2002). The rational of this type of approach is to evaluate a substrate (in this work *U. pinnatifida* exudates) in contact with freshly collected seawater containing the microbial community that will be analyzed. The dilution factor that is used in this approach (0.3 – 0.4) is relatively low with respect to classic culture dilutions, but still allows the response of the communities (growth and degradation) to be easily detected in the short period of time of the experiment. In addition, dilution also reduces grazers pressure, allowing to better profile the bacterial functional capabilities (Carlson et al., 2002). In this experiment, only exudates (and not algae) were added, in order to start with a known concentration of dissolved organic matter, as well as to prevent blades from exudating during the course of this experiment, as this would represent a confounding factor and affect the interpretation of the results.

Exudates are typically composed of polysaccharides, which results in a high C/N ratio, in other words, a nitrogen limitation for microbial growth. A nutrient-amended treatment was therefore included in experiments in order to reduce stoichiometric constraints likely affecting the utilization of exudate-derived-DOC by microbial communities. Nutrients were added in concentrations similar to those found in waters affected by near fishing industries (Torres et al., 2004). Abiotic “Kill” controls were also setup, in order to evaluate whether the observed changes were indeed associated to the action of seawater microbial communities and not due to abiotic factors. Due to the fact that alginate, a major component of macroalgae exudates, is easily decomposed with increased temperature, as well as at pHs lower than 6 or higher than 9, giving a small margin for changing conditions in abiotic controls, formaldehyde was used as inactivating agent (Trevors, 1996).

Briefly, fresh seawater samples were obtained from 5 m depth, near the algal forest. Experimental systems were setup with 200 ml of algal exudate and 100 ml of fresh seawater inoculum, in 300 ml biological oxygen demand (BOD) bottles. Treatments were: i) exudate + fresh seawater (EX) and, ii) exudate + fresh seawater + additional nutrients (EXN, 100 μΜ NH_4_Cl and 10 μΜ KH_2_PO_4_). Incubation controls (INC), consisted in filtered seawater (instead of exudate) + fresh seawater, and abiotic controls (KILL) consisted in exudate + fresh seawater + 1% formaldehyde. Systems were kept in the dark to prevent growth of autotrophic microorganisms, and incubated in these conditions for 5 days. Summarizing, for experiment 2, a total of 32 experimental units were set up. Eight bottles were removed (sacrificed) at each of the four incubation times (0, 24 h, 72 h and 120 h): three replicates corresponding to each of the treatments (EX and EXN), plus one replicate of each of the controls (INC and KILL).

For the analysis of bacterial community structure, experiments were setup as previously described (experiment 2), with the following modifications: (a) exudates were filtered through 0.7 μm GF/F membranes to reduce bacterial load prior to mixing with fresh seawater, (b) a proportion of 210:90 ml exudate:fresh seawater was used, and (c) systems were kept for 7 days, after which samples were retrieved for molecular analyses.

### Metadata and chemical analyses

Temperature, salinity, pH and dissolved oxygen (DO) were measured *in situ* with a multiparameter instrument (Aquacomb). Samples for measuring nutrients were collected in 50 ml acid-washed plastic flasks, and frozen immediately. Nitrate + Nitrite (NO_3_^−^ + NO_2_^−^), phosphate (PO_4_^3−^) and silicate (SiO_3_^4−^) concentrations were determined using a Skalar San Plus autoanalyzer (Skalar Analytical^®^ V.B. 2005a). Ammonium (NH_4_^+^) concentrations were measured manually following the Solórzano’s phenol-hypochlorite method (Strickland and Parsons 1972).

DOC concentrations were measured using a total C analyzer (TOC-L, Shimadzu). Absorbance and fluorescence spectroscopy were applied to characterize the chromophoric and fluorescent fraction of dissolved organic matter (CDOM and FDOM, respectively). For these analyses, 100-ml water samples were serially filtered through glass fiber filters (GF/F, 0.7 μm) and polyethersulfone membranes (PES, 0.2 μm), and stored in precombusted dark bottles at 4°C. The properties of CDOM and FDOM were characterized by absorption and fluorescence spectroscopy (Stedmon and Bro 2008). The absorption spectra of filtered seawater samples were measured in a UV-Visible spectrophotometer (Shimadzu UV1800) and excitation-emission matrices (EEMs) were obtained in a spectrofluorometer (Perkin Elmer LS55). Optical properties of exudates were normalized to DOC concentrations. For details, see **Online Resource 1**.

Total carbohydrates (CH) were measured in exudates by the phenol-sulphuric acid method described in Strickland & Parsons (1972). After the experiments, algae were dried at 45°C to constant weight. DOC and CH were expressed as mg l^−1^ and mg g^−1^ dry weight (dw).

### DOM analysis

Absorbance data were converted to absorption coefficients (a_λ_) following Helms et al. (2008). The water color was calculated as the absorption coefficient at 440 nm (a_440_) and the specific absorbance at 254 nm (SUVA_254_) was calculated as a_254_:DOC and used as an indicator of aromaticity (Weishaar et al. 2003). The spectral slopes for the intervals 275-295 nm (S_275-295_) and 350-400 nm (S_350-400_) and the slope ratio (S_R_= S_275-295_/S_350-400_) were calculated and applied as proxies of molecular weight/size, with higher S values indicating low molecular weight material and/or decreasing aromaticity (Helms et al. 2008; Hansen et al. 2016). In order to characterize the FDOM fraction, the indexes of humification (HIX), autotrophic productivity (BIX), and the fluorescence index (FI, indicative of the relative contribution of terrestrial and microbial sources), were calculated as detailed in **Online Resource 1.** In addition, the FDOM components were obtained applying Parallel Factor Analysis (PARAFAC) to the EEM dataset (Online Resource 1). This analysis performs a mathematical chromatography that allows resolving complex mixture spectra into their pure-component contributions applying three-way modelling (Bro et al., 2010).

For experiment 1, Multiple Linear Correlation analysis and Principal Component Analysis (PCA) were applied to a set of selected response variables (CH and DOC concentration, DOM optical parameters and intensity of the FDOM components). These analyses were carried out using R 3.2.4 (http://www.R-project.org/). PCA was performed with the *pca* function included in the FactoMiner package in R framework. All data were standardized to the variance and centered prior to performing the PCA analysis.

### Bacterial abundance and community alginolytic potential

Bacterial abundance was estimated by colony forming units (CFU) obtained after a 30°C overnight incubation of plates with natural seawater-based culture media (5 g l^−1^tryptone, 1 g l^−1^ yeast extract, 7.5 g l^−1^ agar, prepared in filtered and UV-treated natural seawater). The potential of the microbial community to degrade alginate was tested *in vivo* in liquid assays consisting of 600 μl seawater samples and 200 μl of 0.2% sodium alginate, incubated overnight at room temperature. The alginolytic potential of the sample was estimated by the increase in absorbance at 235 nm between the initial time (T=0) and 24 h (T=24). This increase is indicative of the formation of a double bond between C-4 and C-5 at the new non-reducing end of the polysaccharide by the β-elimination reaction catalyzed by the alginate lyase enzyme (Cao et al. 2007).

### Bacterial community structure

For molecular analysis of bacterial communities, 20 ml samples were obtained from experiment 2 incubations, centrifuged 30 min at 20,000 *g* in a Sorvall Refrigerated Superspeed Centrifuge (RC-5C), and resuspended in 1 ml of 50 mM TrisHCl buffer pH 8.0. DNA was extracted using a method modified from Bostrom and collaborators (Boström et al. 2004). Briefly, cell pellets were resuspended in lysis buffer (400 mMNaCl, 750 mM sucrose, 200 mM EDTA, 50 mMTrisHCl buffer pH 8), and lysozyme was added to a final concentration of 1 mg/ml. The suspensions were kept at 37°C for 30 min. Then, sodium dodecyl sulphate (SDS, final concentration of 1% w/v) and Proteinase K (to a final concentration of 0.1 mg/ml) were added, and samples were kept at 55°C overnight. After the lysis, 0.1 volume of 3M sodium acetate and one volume of isopropanol were added, and the samples were incubated overnight at −20°C to precipitate the DNA. Samples were centrifuged in a microcentrifuge for 30 min at maximum speed (~16,000 *g*), washed with 70% ethanol prepared in molecular-grade distilled water, and dried for 10 min at 50°C. DNA pellets were resuspended in 50 μl of 10 mM TrisHCl buffer pH 8 prepared in molecular-grade distilled water.

Changes in microbial community structure were analyzed by large-scale 16S rRNA gene amplicon sequencing, with universal primers spanning the V4 hypervariable region (515F (Parada et al. 2016): 5’GTGYCAGCMGCCGCGGTAA3’ –806R (Apprill et al. 2015), 5’GGACTACNVGGGTWTCTAAT3’, forward-barcoded, Earth Microbiome Project, updated for increased Archaea and SAR11 coverage, http://www.earthmicrobiome.org/protocols-and-standards/16s/). Sequences were processed with QIIME2 (Bolyen et al. 2019) using DADA2 to identify amplicon sequence variants (ASV) (Callahan et al. 2016, 2017). ASVs were classified in QIIME2 using Greengenes classifier (https://data.qiime2.org/2019.1/common/gg-13-8-99-515-806-nb-classifier.qza). For details of sequence processing, see **Online Resource 2**. The resulting feature table was analyzed in R environment with packages *phyloseq* (McMurdie and Holmes 2013)*, vegan* (Oksanen et al. 2019), and *microbiome* (Lahti et al., 2017). The software *picrust* (Douglas et al. 2019) was used to estimate the metabolic functions from 16S rRNA gene amplicon sequence data, using KeggOrthology (KO) terms and pathways (Kanehisa and Goto 2000). *Picrust* infers the gene families likely to be present in a sample, based on the genomic information available for the taxonomic assignment of each ASV. The *picrust2* version used in this study is compatible with denoising methods, wrapped into QIIME2.

### Statistical analyses

One tail Welch’s Two Sample t-test was performed on CH and DOC data from experiment 1 to test the null hypothesis that true difference in means between *U. pinnatifida* and control incubations is less or equal to 0. A test for association/correlation between paired samples, using Pearson’s product-moment correlation index, was performed between selected chemical (DOC, CH) and microbiological (abundance, alginolytic potential) variables, for experiment 1. All analyses were performed in R environment.

## Results

### Characterization of *U. pinnatifida* exudates

During experiment 1, algae released exudates that were yellowish and viscous, dense like mucus. No precipitate could be observed. Algae released an average of 1.6 ± 0.8 mg C g^−1^ biomass day^−1^ into seawater, with more than half (0.9 ± 0.5 mg g^−1^ day^−1^) corresponding to carbohydrates. Carbohydrates increased rapidly, as well as DOC (**Table 1**). The characterization of the fluorescent fraction of the DOC through parallel factor analysis (PARAFAC) based on the excitation-emission matrices (EEM) collected from experiment 1 allowed decomposing the mixture of compounds into underlying individual fluorophores constituting collectively the FDOM fingerprinting (Bro et al., 2010). Three fluorescent components (C) were modelled: C1 and C2 (a combination of peaks A + M and A + C, respectively) attributable to humic compounds; and C3 (peak T), which is a “non-humic” component (Coble 1996; Stedmon et al. 2003). The comparison of these components against the worldwide database Openfluor confirmed the correspondence of C1 and C2 with humic-like compounds (Yamashita et al. 2010; Cawley et al. 2012; Catalá et al. 2015) and associated C3 with protein-like, aliphatic compounds likely derived from recent microbial production (Maie et al. 2007; Kellerman et al. 2015; Osburn and Bianchi 2016) (for details see **Online Resource 1**)..

**Table 1.**
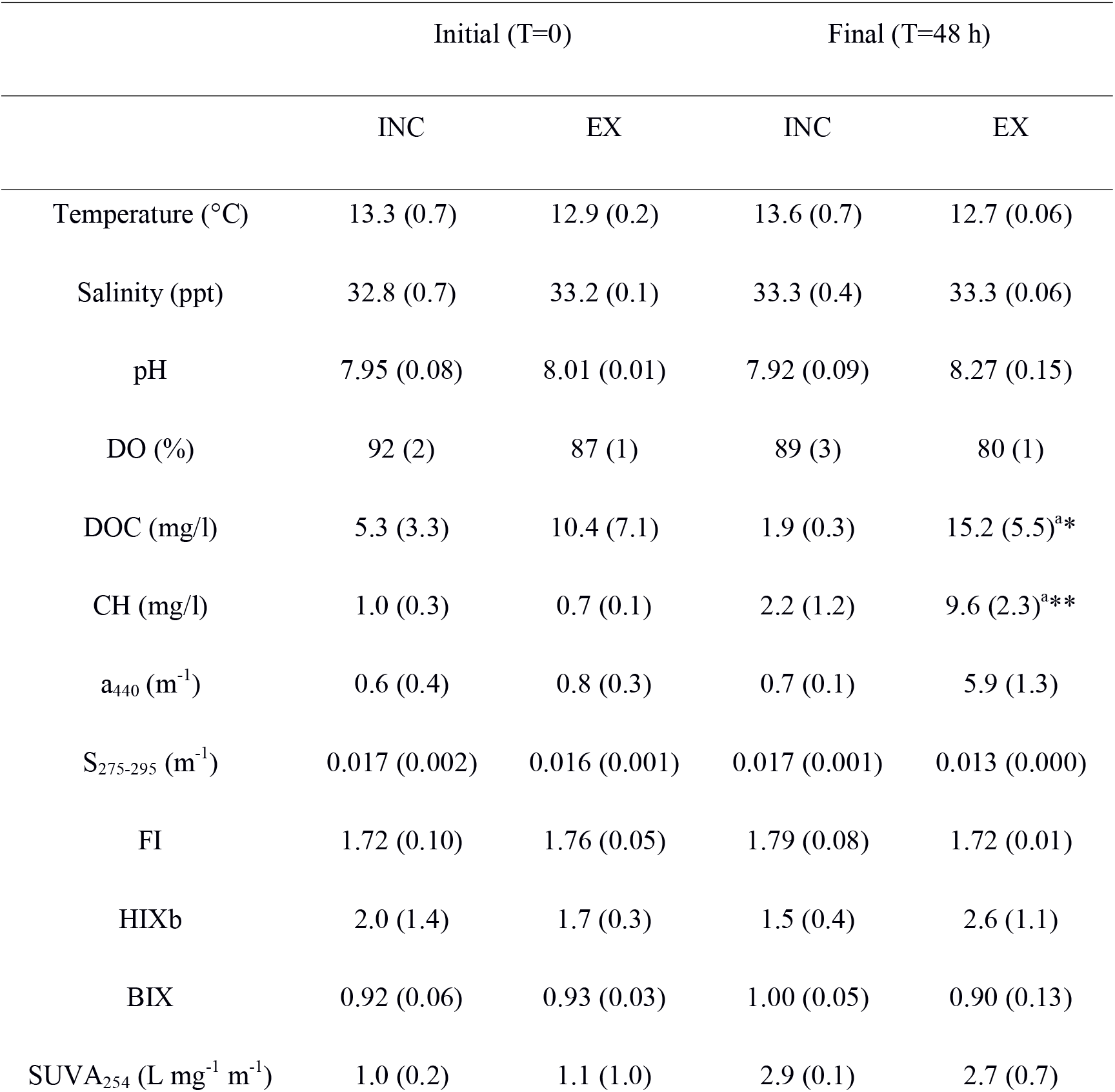
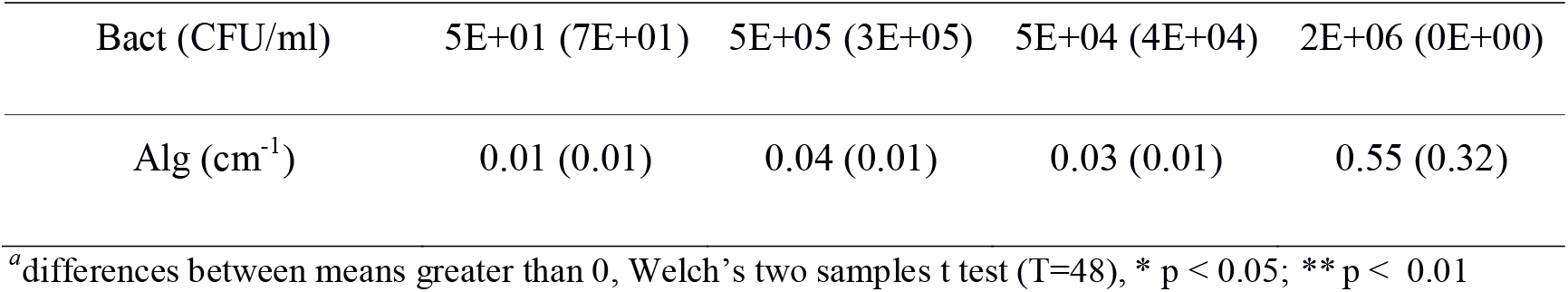
Changes in physicochemical and microbiological parameters in *U. pinnatifida*-seawater incubations. DO: Dissolved oxygen. DOC: dissolved organic carbon. CH: Total carbohydrates concentration. a_440_: absorbance at 440 nm. S_275-295_: spectral slope calculated for the decrease in absorbance between 275 and 295 nm. FI: fluorescence index. BIX: biological index. HIX: humification index. SUVA_254_: Absorbance at 254 nm:DOC. Bact: colony forming units (CFU) in heterotrophic plate counts (estimation of aerobic bacterial abundance). Alg: alginolytic potential, measured as alginate lyase activity in an *in vivo* liquid assays. EX: incubation systems with *U. pinnatifida*, INC: control systems with only seawater. Values are means (± 1 standard deviation).

|  | Initial (T=0) |  | Final (T=48 h) |  |
| --- | --- | --- | --- | --- |
|  | INC | EX | INC | EX |
| Temperature (°C) | 13.3 (0.7) | 12.9 (0.2) | 13.6 (0.7) | 12.7 (0.06) |
| Salinity (ppt) | 32.8 (0.7) | 33.2 (0.1) | 33.3 (0.4) | 33.3 (0.06) |
| pH | 7.95 (0.08) | 8.01 (0.01) | 7.92 (0.09) | 8.27 (0.15) |
| DO (%) | 92 (2) | 87 (1) | 89 (3) | 80 (1) |
| DOC (mg/l) | 5.3 (3.3) | 10.4 (7.1) | 1.9 (0.3) | 15.2 (5.5)<sup>a*</sup> |
| CH (mg/l) | 1.0 (0.3) | 0.7 (0.1) | 2.2 (1.2) | 9.6 (2.3)<sup>a**</sup> |
| $a_{440}$ (m<sup>-1</sup>) | 0.6 (0.4) | 0.8 (0.3) | 0.7 (0.1) | 5.9 (1.3) |
| $S_{275-295}$ (m<sup>-1</sup>) | 0.017 (0.002) | 0.016 (0.001) | 0.017 (0.001) | 0.013 (0.000) |
| FI | 1.72 (0.10) | 1.76 (0.05) | 1.79 (0.08) | 1.72 (0.01) |
| HIXb | 2.0 (1.4) | 1.7 (0.3) | 1.5 (0.4) | 2.6 (1.1) |
| BIX | 0.92 (0.06) | 0.93 (0.03) | 1.00 (0.05) | 0.90 (0.13) |
| $SUVA_{254}$ (L mg<sup>-1</sup> m<sup>-1</sup>) | 1.0 (0.2) | 1.1 (1.0) | 2.9 (0.1) | 2.7 (0.7) |
| Bact (CFU/ml) | 5E+01 (7E+01) | 5E+05 (3E+05) | 5E+04 (4E+04) | 2E+06 (0E+00) |
| Alg (cm<sup>-1</sup>) | 0.01 (0.01) | 0.04 (0.01) | 0.03 (0.01) | 0.55 (0.32) |

<sup>a</sup>differences between means greater than 0, Welch's two samples t test (T=48), \* p < 0.05; \*\* p < 0.01

Both an increase in bacterial abundance and alginolytic potential (the capability of the communities of degrading alginates, evaluated in overnight incubations) accompanied exudate release (**Table 1**), and were directly correlated with CH concentration (Pearson’s r = 0.85 and r = 0.94, for bacterial abundance and alginolytic potential respectively, p<0.05), while the alginolytic potential was also correlated to total DOC (r = 0.71, p < 0.05).

In order to better display the relationships in selected response variables measured in the different treatments, multiple correlation analysis (MC) and principal component analysis (PCA) were applied to the resulting data. For experiment 1, selected response variables included DOC and CH concentrations, the optical parameters a_440_, SUVA_254_; S_275-295_, the fluorescence indexes HIX, BIX, FI and the fluorescent components C1 (peaks A+M), C2 (peaks A+C), C3 (peak T). The results are shown in **Figure 1** (PCA) and **Online Resource 3**.

**Fig. 1.**
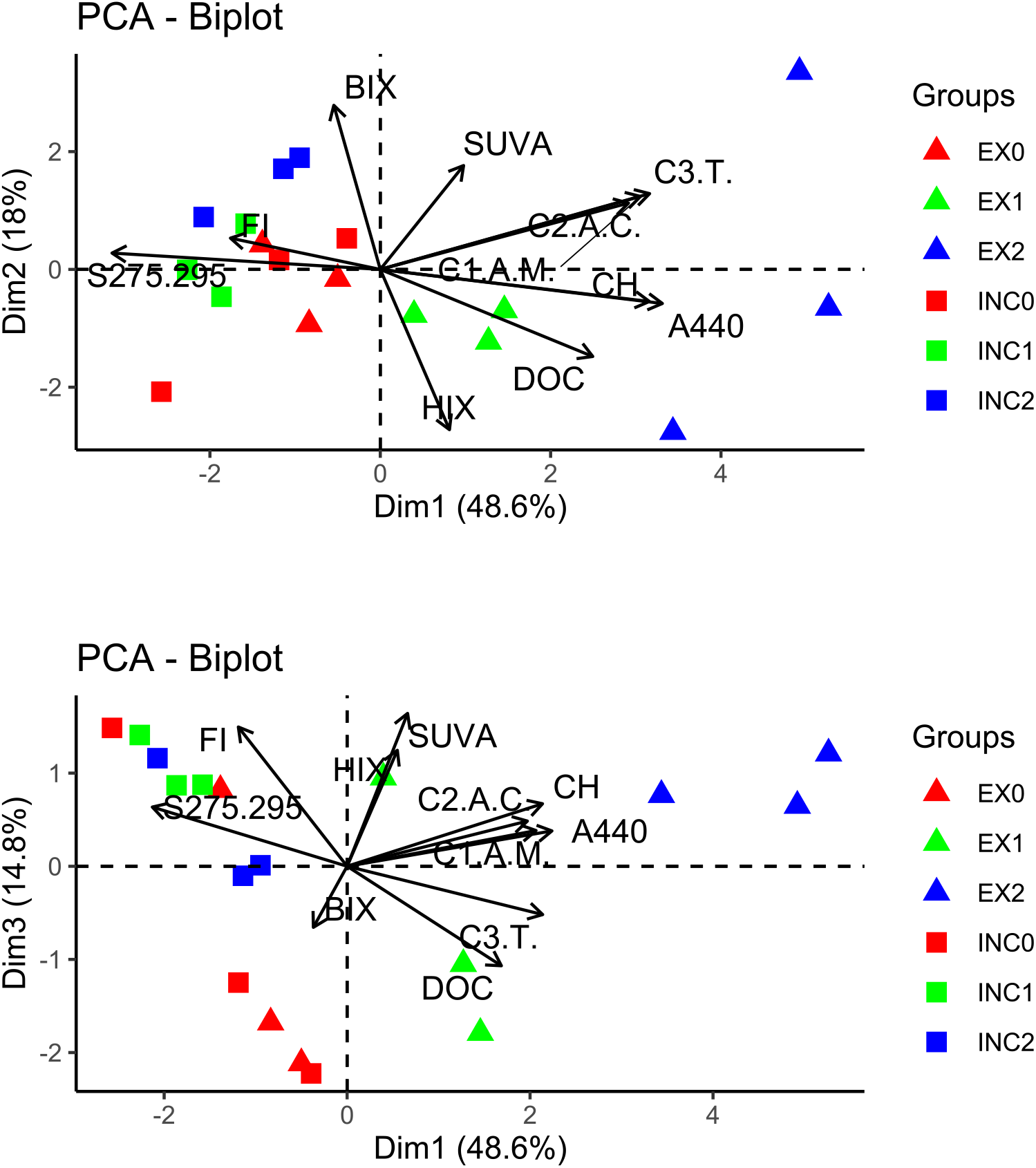
Principal Component analysis, performed to the set of response variables measured in experiment 1. PCA plots: PC1 vs PC2 (upper panel) and PC1 vs PC3 (lower panel). The samples are named as condition (INC or EX) followed by day of sampling (0, 1 or 2)

In addition to increasing DOC and CH concentrations, *U. pinnatifida* exudates concomitantly increased the seawater color (a_440_) and the intensity of the three DOM fluorescent components C1, C2 and C3 in seawater (Online Resource 3). Throughout the incubation with *U. pinnatifida*, the molecular weight/ size and the aromaticity of the DOC increased continuously, as indicated by the negative correlation observed between the S_275-295_ (inversely related with DOM molecular weight/size and degree of aromaticity), and the three fluorescent components, the water color and the DOC and CH concentrations (**Online Resource 3**). The negative correlation between DOC concentration and the FI and the component C3 (peak T, associated to microbial activity) indicated bacterial consumption on the non-humic and more biolabile DOC fraction. The negative correlation between the indexes BIX and HIX suggests that the biodegradation of the labile DOC fraction led to an increase in DOM humification (**Online Resource 3**).

As regards the PCA, the first three principal components explained 84% of the variance of the dataset, with PC1 explaining 53%, PC2 19% and PC3 12.2% of the variance. The PC1 correlated positively with the three fluorescent DOM components, the a_440_ and the DOC and CH concentrations, and negatively with the S_275-295_ (**Online Resource 3**). Along the PC1, three groups were apparent: controls at all times with exudate treatments at day 0, day1, and day 2 (**Figure 1, upper panel**). Therefore, PC1 can be related to the process of algae exudation in the experimental systems. On the other hand, PC2 correlated positively with the BIX and negatively with the HIX, and PC3 with the FI (**Online Resource 3**), variables related to microbial processing of the recently released organic matter. These components also revealed a strong variation among replicates in the different conditions (**Figure 1, lower panel**).

Overall, these results show that the incubation of *U. pinnatifida* in seawater increased the DOC concentration shifting also DOM quality. In particular, during the incubation in experiment 1, algal exudates increased the CH concentration, humic substances (included in the fluorescent components C1 and C2) and non-humic components (represented in the C3). In this last component, a contribution to the DOM pool of the microbial community attached to the algae cannot be discarded.

### Further processing of *U.pinnatifida* exudates by seawater microbial communities

In experiment 2, fresh seawater microbial communities from PE were exposed to *U. pinnatifida* exudates obtained in experiment 1, with the aim of analyzing the response of bacterioplankton to these exudates, as well as the effects of microbial processing on the DOM pool. In addition to EX treatment, EXN systems were setup, in which the macronutrients NH_4_ and PO_4_ were provided at non-limiting concentrations [> 100 and >10 μM, respectively, concentrations similar to those found in waters affected by near fishing industries (Torres et al. 2004)]. For details see **Online Resource 4**.

In addition to a decrease in DO and aerobic heterotrophic bacterial abundance, only moderate changes in DOC or CH concentrations were observed in the course of experiment 2 (**Figure 2**). Differents in initial DOC and CH were already present (Figure 2, t=0), and were carried throughout the course of the experiment. This is likely chemical variability associated with the colloidal nature of the exudates, which is a source of variation among replicates. Exudates were very viscous and difficult to filter (as mentioned in the first section of results) so this probably reflects the natural conditions experienced by seawater bacterial communities near the algal banks. In contrast to the quantitative results, which were not conclusive, the quality of DOM analyzed in terms of the absorbance and fluorescence properties (Helms et al. 2008; Hansen et al. 2016) evidenced progress in diagenesis throughout the experiment. For example, an increasing tendency in the SUVA_254_ was observed in the EX and EXN treatments with respect to the controls, indicating progressive humification along the incubation (**Figure 3A**). The S_275-295_ remained low with respect to controls **(Figure 3B**), which may relate to an increase in the relative contribution of recalcitrant material, such as aromatic compounds, due to the biodegradation of the more labile DOM fractions.

**Fig. 2.**
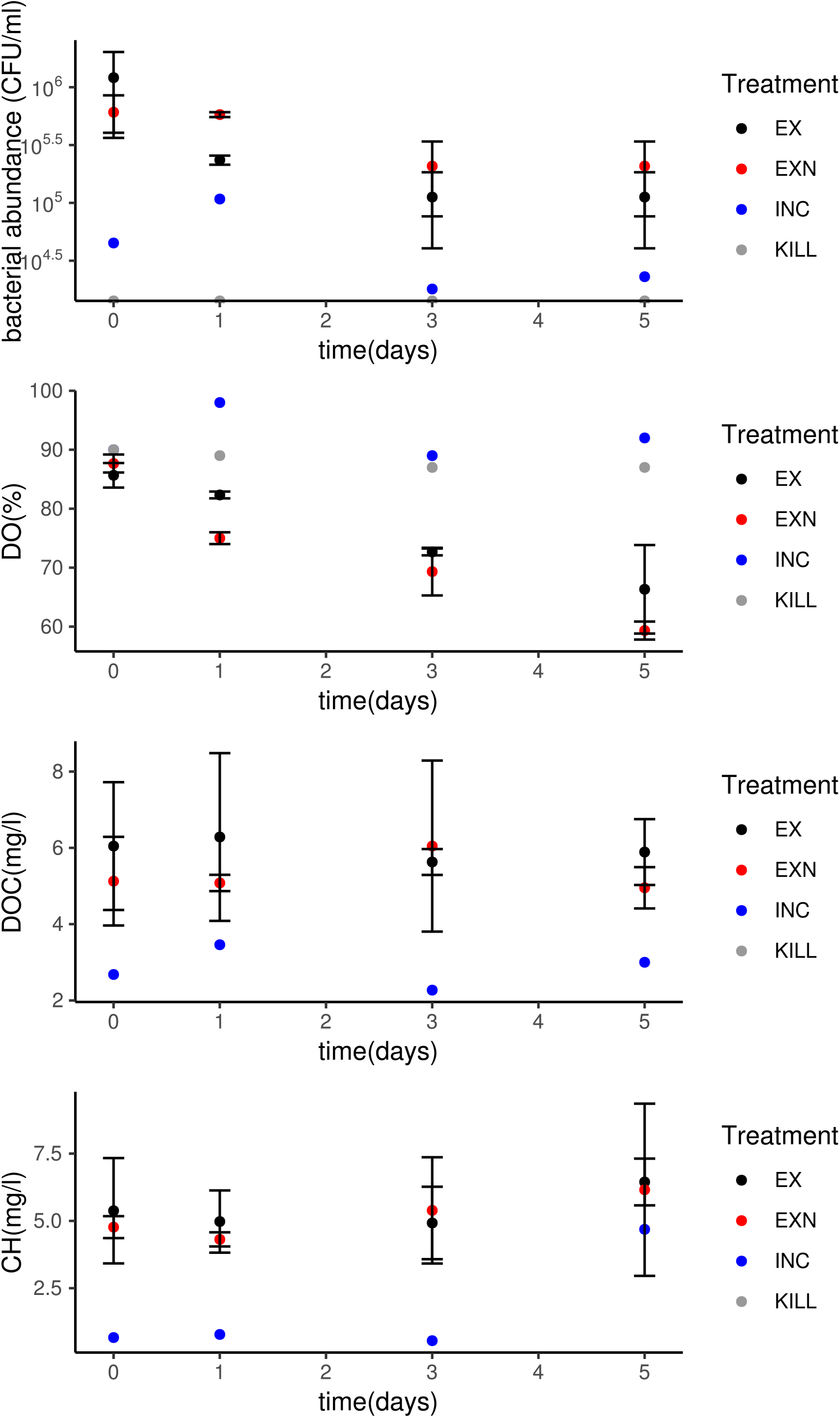
Changes in physicochemical and microbial parameters in the different treatments applied in experiment 2 (processing of *U. pinnatifida* exudates by the seawater microbial community). EX: exudate + fresh seawater, EXN: exudate + fresh seawater + additional nutrients, INC: filtered sterile seawater + fresh seawater (control), KILL: exudate + fresh seawater + 1% formaldehyde, DO: dissolved oxygen, DOC: dissolved organic carbon, CH: carbohydrates content

**Fig. 3.**
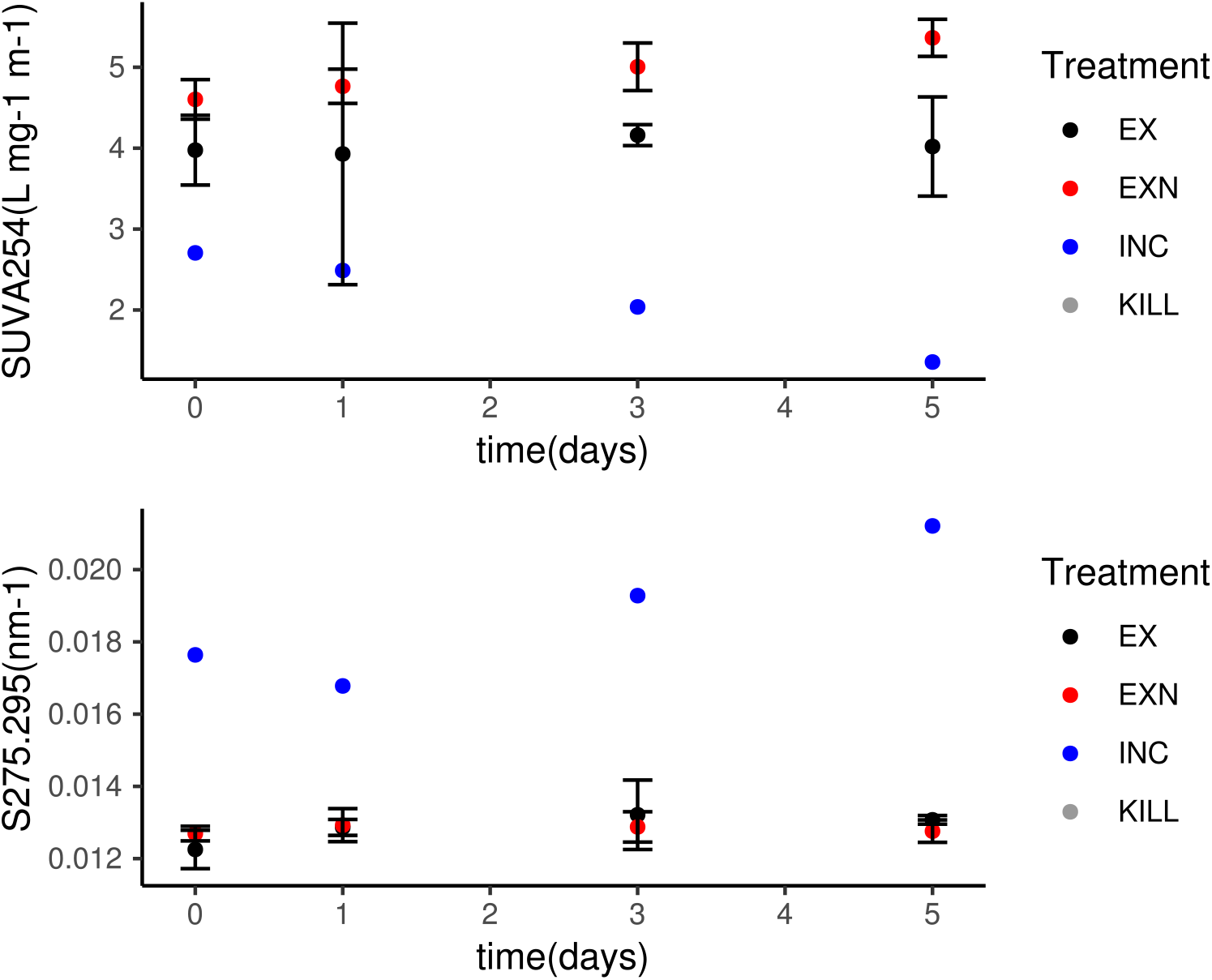
Changes in the optical properties of exudate-derived DOM in the experimental systems, along incubation of PE seawater with *U. pinnatifida* exudates (experiment 2). EX: exudate + fresh seawater, EXN: exudate + fresh seawater + additional nutrients, INC: filtered sterile seawater + fresh seawater (control), KILL: exudate + fresh seawater + 1% formaldehyde. SUVA_254_: Absorbance at 254:DOC, S_275-295_: spectral slope calculated for the decrease in absorbance between 275 and 295 nm.

The PCA performed to experiment 2 data showed that three principal components explained 79% of the variance, with PC1 explaining 53%, PC2 17.6% and PC3 8.5% of the variance. The PC1, basically related to differences among treatments (INC controls and exudates incubations, **Figure 4**), correlated positively with CH, DOC, a_440_, SUVA_254_, FI, and the two fluorescent DOM components C1(A+M), C2 (A+C), and negatively with the S_275-295_ and BIX (**Online Resource 5**). On the other hand, PC2 correlated positively with FDOM component C3(T) and negatively with the HIX (**Online Resource 5**), results that are mainly explained by differences between EX and EXN treatments, and are related to an increase in component C3 by diagenesis in the presence of added nutrients. The component PC3 showed correlation with aerobic bacterial abundance (CFU/ml, **Online Resource 5**).

**Fig. 4.**
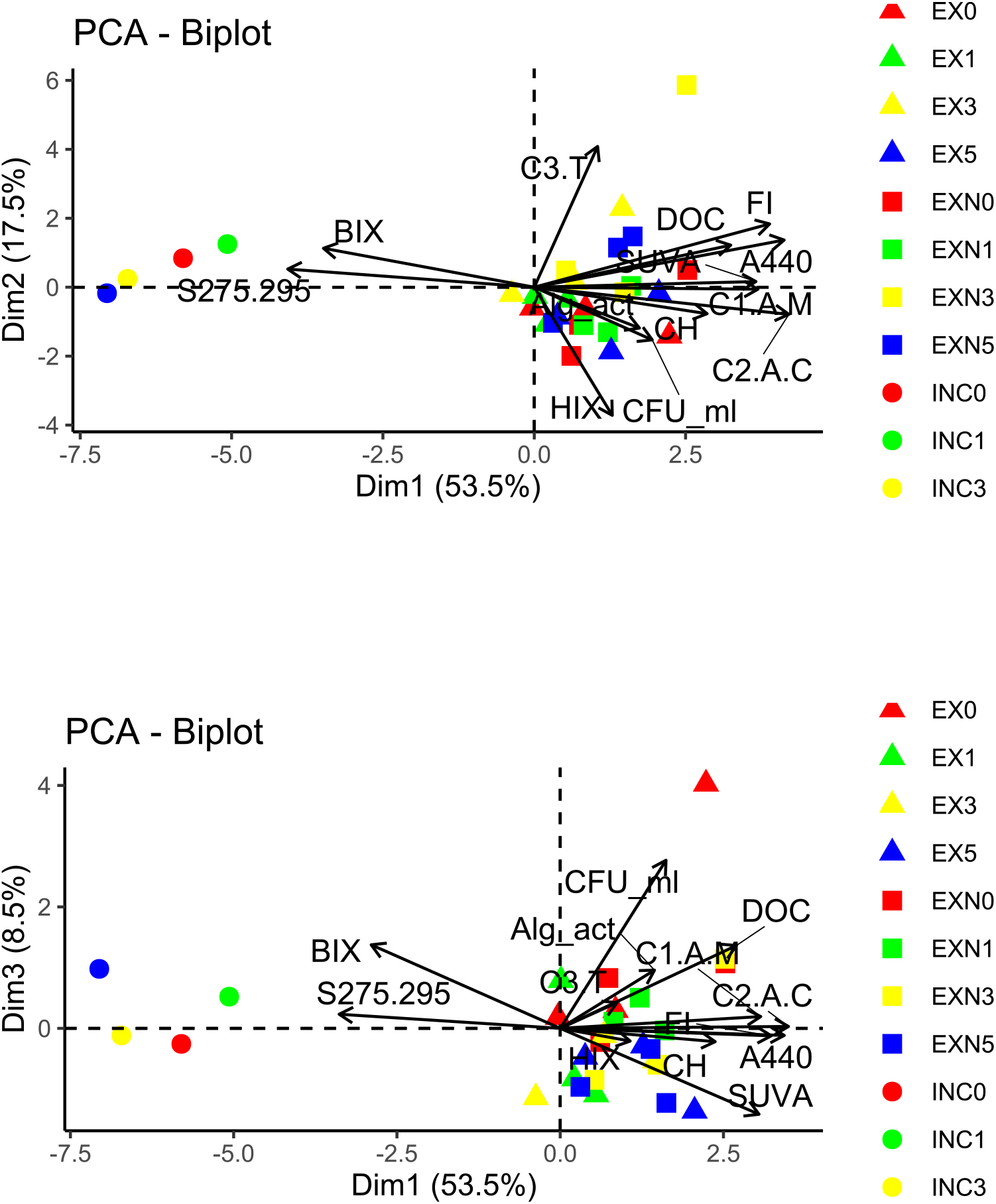
Principal Component analysis, performed to the set of response variables measured in experiment 2. PCA plots: PC1 vs PC2 (upper panel) and PC1 vs PC3 (lower panel). The samples are named as condition (INC or EX) followed by day of sampling (0, 1 or 2)

Overall, the results show that exudates undergo a very rapid biodegradation characterized by labile carbon depletion in the first days. The decrease in aerobic bacterial cell numbers observed was probably due to the depletion of oxygen and labile carbon sources. Also, while the dilution used in the experiment probably limited grazing (Carlson et al., 2002), we cannot exclude the possibility of cell death due to predation by microzooplankton. Bacterial cell death may have also contributed to increase the DOC concentration and shift the DOM quality from the prevalence of relatively labile (algal carbohydrates) to more recalcitrant algal but also bacterial-derived DOM (Ogawa et al. 2001; Lechtenfeld et al. 2015).

### Shifts in seawater bacterial communities after exudate enrichment

In order to gain a better insight into the effect of *U. pinnatifida*-derived DOM on seawater bacterial communities, large-scale 16S rRNA gene amplicon sequencing was carried out from experimental systems consisting of exudates and seawater from PE site, with and without added nutrients (EXN and EX treatments), in a similar way as described above. Experimental systems were incubated for 7 days at 14°C, and initial and final samples were collected for DNA extraction (**Table 2, Online Resource 4**).

**Table 2.**
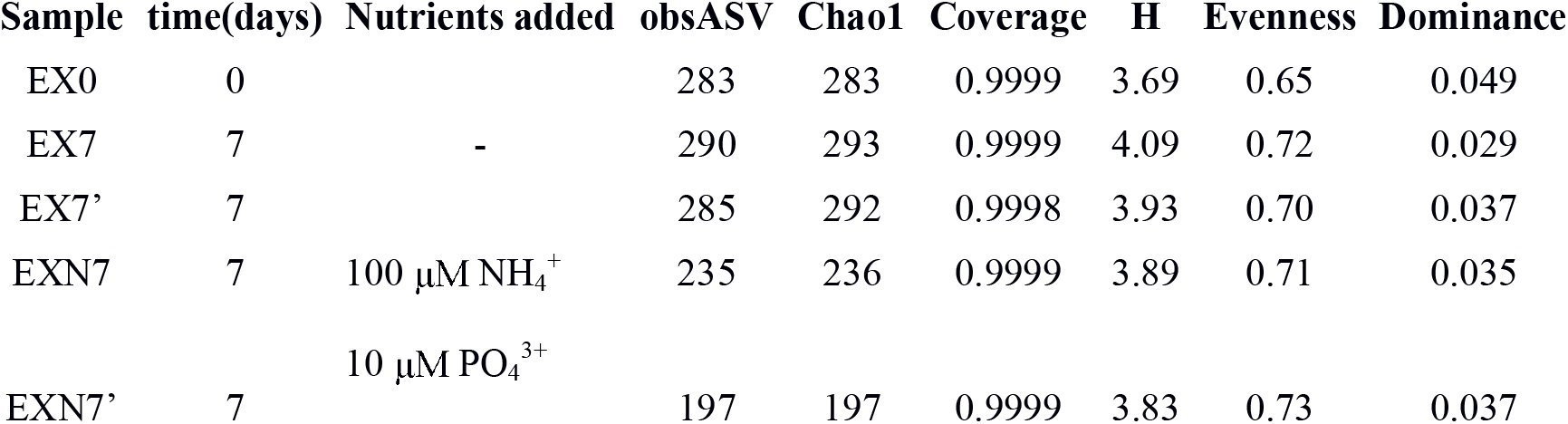
Alpha diversity estimators for microbial communities enriched after exposure to *U. pinnatifida* exudates. Samples were rarefied to ~60,000 sequences per sample, according to the sample that yielded less sequences. ObsASV: observed amplicon sequence variants. Chao1: Chao’s richness estimator. Coverage: Good’s coverage index. H: Shannon’s diversity index. Evenness: Pielou’s index. Dominance: Simpson’s dominance index.

After preprocessing the sequences, a total of 492 amplicon sequence variants (ASV) were observed to be present in the community, with ~ 200 ASV per individual sample (**Table 2**). Coverage values (Good’s coverage, an indicator of how completely the richness of the community has been sampled) were higher than 99.99% for all samples, indicating that the sequencing depth of 60,000 sequences per sample was very efficient in recovering the bacterial diversity present in these systems. Although richness (i.e. number of estimated variants, by Chao1 index) was lower in the samples from EXN treatment, estimators taking into account the relative abundance (such as Shannon’s diversity index or Pielou’s evenness) showed an increasing tendency in all experimental systems after 7 days. The opposite was observed for dominance (**Table 2**).

### Sample time(days) Nutrients added obsASV Chao1 Coverage H Evenness Dominance

Community similarity analysis showed a clear distinction between communities at initial conditions (EX, T=0) and after 7 days in both experiments (EX, exudate-exposure and EXN, exudate and nutrients exposure, T=7) (**Figure 5a**). This result indicates changes in seawater bacterial populations after various days of contact with *U. pinnatifida* exudates. Such effects could be either direct by the exposure to its carbon sources or indirect due to changes in fundamental variables such as dissolved oxygen, which led to shifts in community structure. Differences in community structure between nutrient-amended and not amended treatments indicate that nutrients have a role in shaping microbial assemblages, e.g. by selecting populations with different nutrient affinities and/or uptake efficiencies. Initially, bacterial communities were dominated by members of the Gammaproteobacteria, mainly belonging to the orders Alteromonadales, Vibrionales, and Oceanospirillales. Alphaproteobacteria of the order Rhodobacterales were also abundant (**Figure 5b**). This structure is very similar to that commonly observed in natural coastal bacterioplankton (Taylor and Cunliffe 2017), but also in seawater that has been exposed to algal exudates (Nelson et al. 2013). These microorganisms are probably members of the natural seawater bacterial communities of Punta Este site, which are periodically exposed to algal exudates. In our experimental set up, however, we cannot rule out that some of them could have passed with the exudate through the 0.7 μm filter.

**Figure 5.**
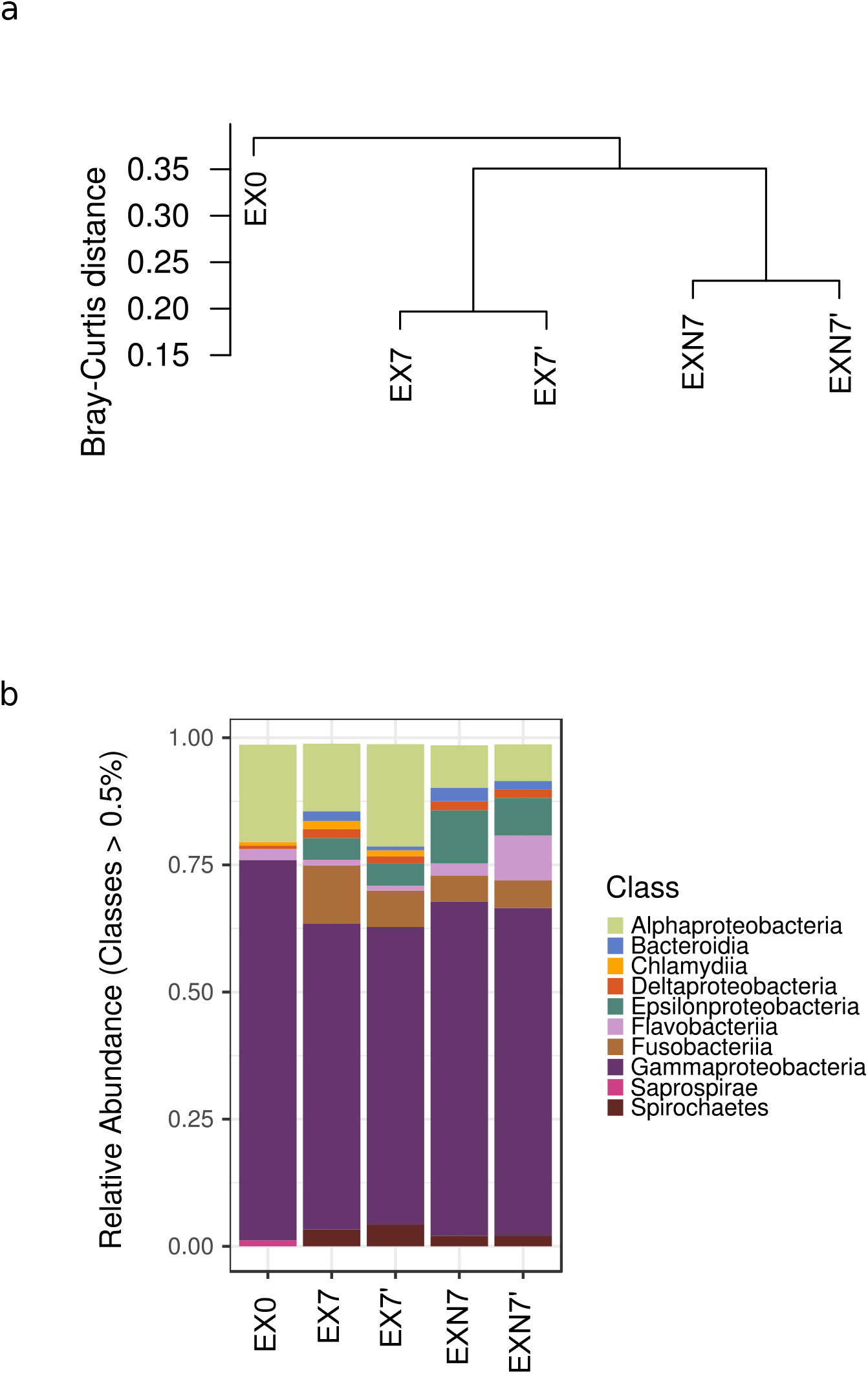
Bacterial community structure of *U. pinnatifida* exudate-seawater experimental systems, based on 16S rRNA gene amplicon sequencing. **A-** Hierarchical clustering (UPGMA) of samples at the ASV level. Bray-Curtis dissimilarity was used as distance measure. B-Taxonomic assignment of ASVs at the Order level. Sample EX0: initial conditions. Samples EX7 and EX7’: duplicate exudate enriched experiments after 7 days of incubation. EXN7, EXN7’: duplicate exudate enriched experiments with added nutrients

We analyzed community changes in more detail, identifying the ASVs that showed positive or negative differences in abundance with respect to the initial conditions. Some of the initially dominant populations were present at lower abundances at the end of the experiment **(Figure 6**, bubbles with y-values lower than zero), while others increased their abundances (**Figure 6,** positive y-values). We considered as “differentially abundant ASVs” those shifting in more than 1% (either negative or positive) in both replicates with respect to the initial time (for details of all abundances as well as taxonomic assignments of ASVs see **Online Resource 6**).

**Fig. 6.**
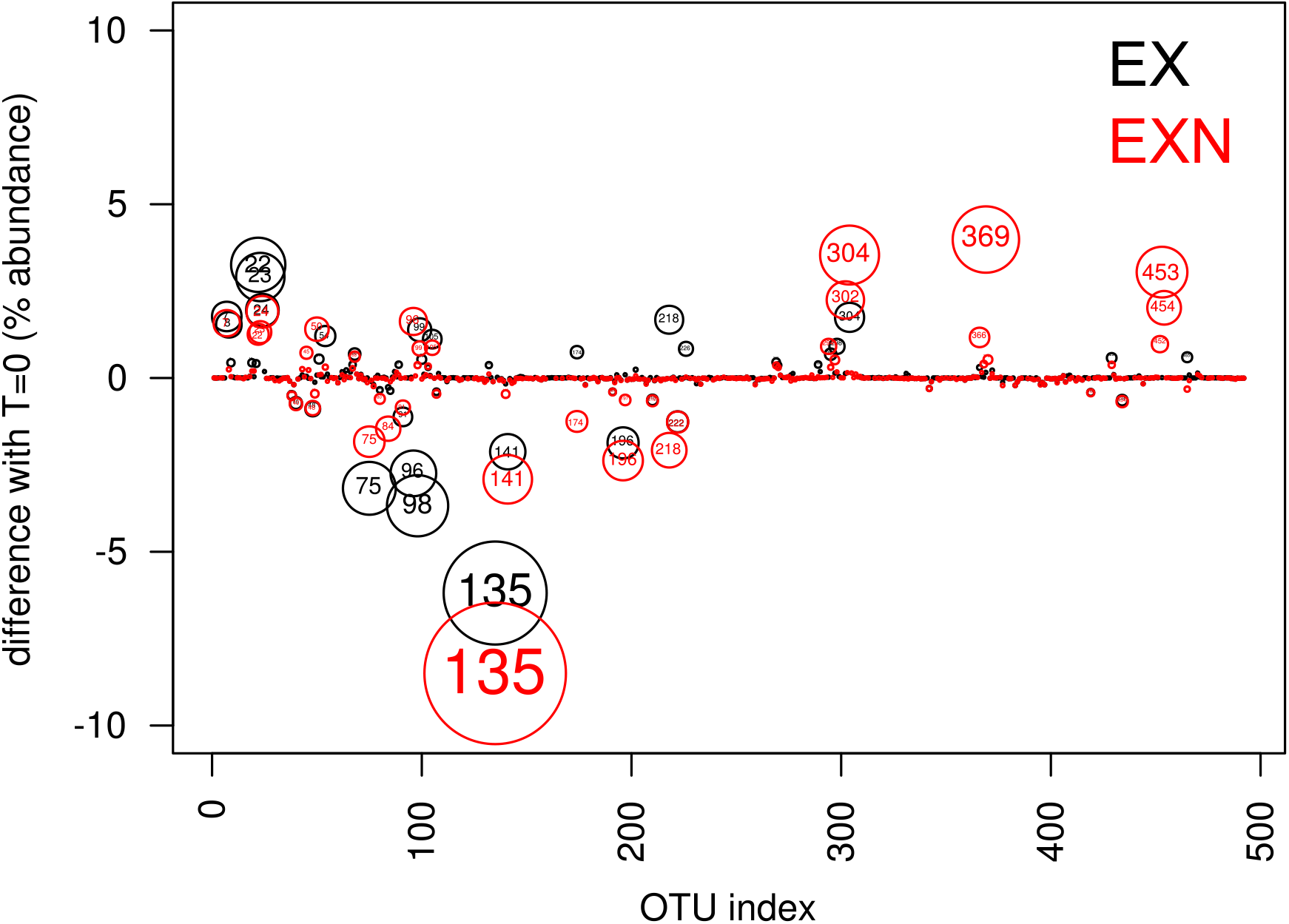
Differentially abundant ASVs in experimental systems containing seawater enriched with *U. pinnatifida* exudates. Values are average differences between % abundance in each condition (EX: exudate, EXN: exudate plus nutrients) with respect to initial conditions. The size of the bubbles is proportional to the difference in abundances for each ASV

Of the variants that were initially dominant, nine decreased after incubation: ASVs #75, #84, #91, #96, #98, #135, #141, #196, and #222. These ASVs were classified as members of the Gammaproteobacteria: *Alteromonas* and *Psychromonas* (Alteromonadales), *Vibrio* (Vibrionales), as well as Alphaproteobacteria (unclassified Rhodobacteraceae) (**Table 3**). On the other hand, 14 ASVs showed an increase in abundance: #7, #8, #22, #23, #24, #50, #54, #99, #105, #302, #304, #369, #453, and #454 (**Table 3**). Seven of these (#7, #8, #22, #23, #24, #105 and #304) increased with 1% difference in both EX and EXN. These were assigned as belonging to *Spirochaeta* (Spirochaetes)*, Propionigenium* (Fusobacteria), *Shewanella* (Gammaproteobacteria, Alteromonadales) and *Arcobacter* (Epsilonproteobacteria, Campylobacterales). The ASVs belonging to *Vibrio* showed a variant-specific behaviour: some decreased with exudates exposure but increased with exudate plus nutrients, while others increased in both conditions (**Table 3**). Notably, the differentially abundant ASVs accounted for an important fraction of the reads, indicating a drastic shift in these communities (**Table 3**).

**Table 3.**
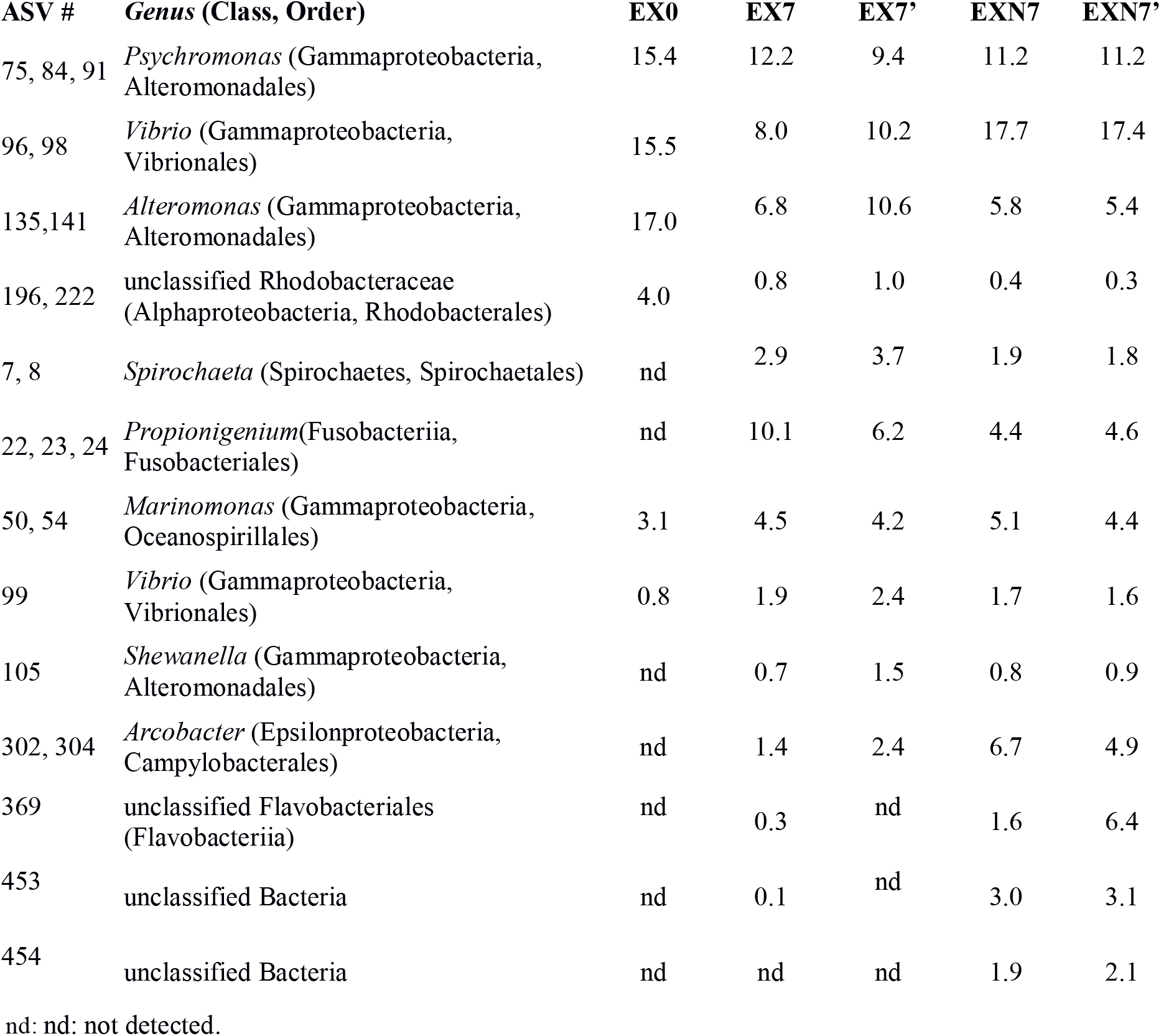
ASVs with differences in abundance in the experimental systems, with respect to initial conditions. Values are percent abundances out of total 16S rRNA gene amplicon sequences. EX0: initial conditions. EX1 and EX2: exudate enriched experiments, after 7 days. EXN1, EXN2: exudate enriched experiments with added nutrients, after 7 days.

Some variants were clearly related to nutrient addition (EXN). This was the case of the ASV #96 and #98 (classified as *Vibrio*), and of #369 (belonging to the Flavobacteriales) which reached high relative abundances in EXN but was very low in all other conditions. This variant could not be assigned to the genus level, indicating that it could constitute a novel group. Two variants that shared high identity values (99.6%), #453 and #454, could only be classified as “Bacteria”. When analyzed by blastn search, these sequences were most closely related (although with only 84% identity), to the 16S rRNA gene from *Kiritimatiella glycovorans*, a genus assigned to a novel phylum from the PVC superphylum (Kiritimatiellaeota, (Spring et al. 2016)).

Given the important changes observed in community structure, we utilized phylogenetic information (16S rRNA gene amplicon sequence data) to infer changes in the metabolic potential associated to the shifts observed after exposure to *U. pinnatifida* exudates. This analysis was performed using the *picrust* program (Douglas et al. 2019), which analyzes the genomic information available in public databases for each genus detected in the sample by amplicon sequencing. The gene family information (KO terms) was inferred for each ASV and was then assigned to the corresponding KEGG pathways. Pathways that were differentially abundant in the 7-day experiment samples with respect to initial condition were identified (**Figure 7)**.

**Fig. 7.**
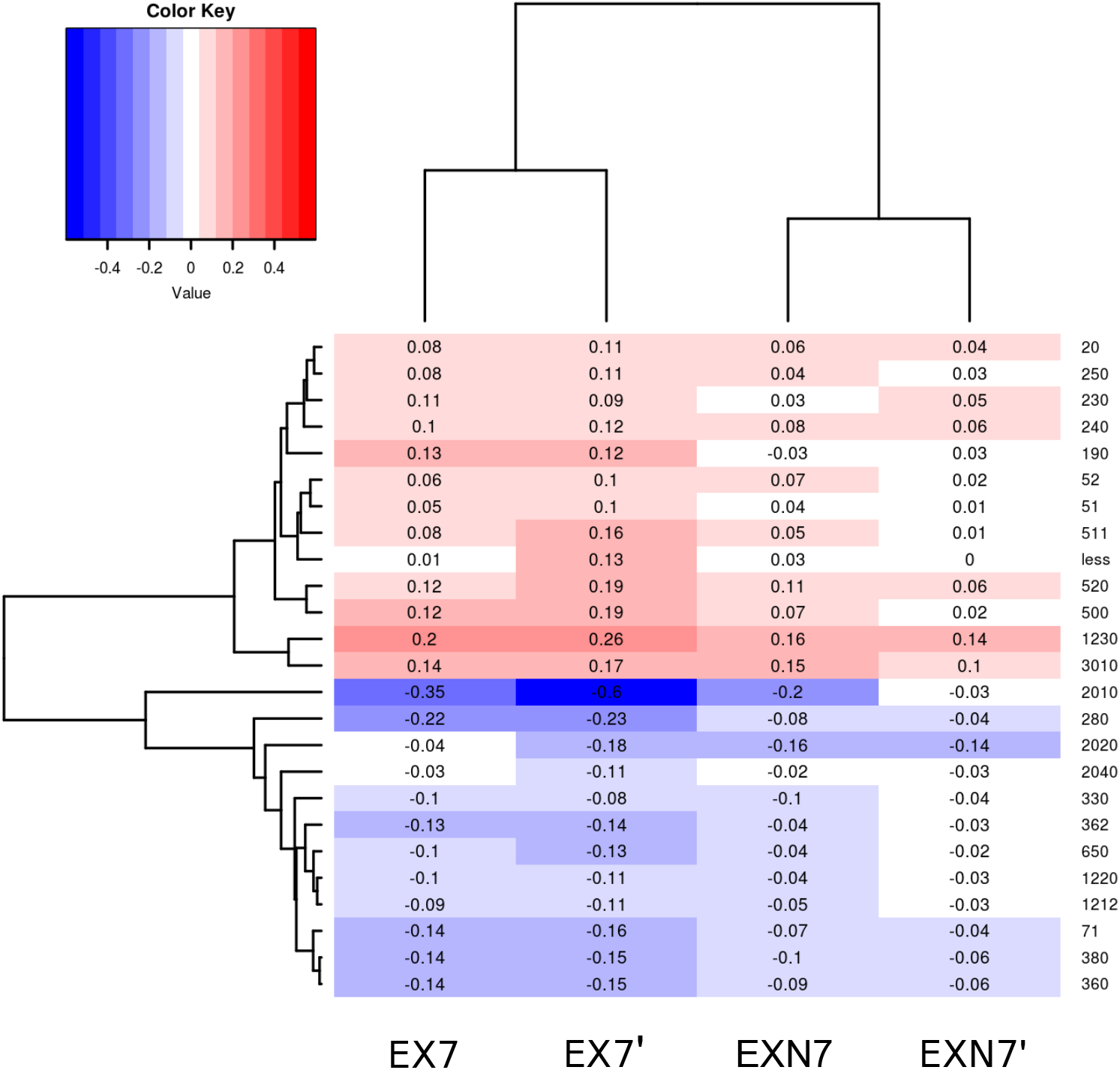
Differentially abundant pathways in experiments of seawater exposure to exudates from *U. pinnatifida,* as inferred from amplicon sequence data using *picrust2*. Values are percent differences with respect to the initial conditions (EX0). Only the pathways displaying differences of more than 0.1% with respect to EX0 in at least one of the samples are shown. Red: pathways with estimated abundances higher than EX0, *i.e.* increased after exudate or exudate-nutrients addition. Blue: pathways that decreased. Pathway keys: 00020 Citrate cycle (TCA cycle), 00250 Alanine, aspartate and glutamate metabolism, 00230 Purine metabolism, 00240 Pyrimidine metabolism, 00190 Oxidative phosphorylation, 00052 Galactose metabolism, 00051 Fructose and mannose metabolism, 00511 Other glycan degradation, 00520 Amino sugar and nucleotide sugar metabolism, 00500 Starch and sucrose metabolism, 01230 Biosynthesis of amino acids, 03010 Ribosome, 02010 ABC transporters, 00280 Valine, leucine and isoleucine degradation, 02020 Two-component system, 02040 Flagellar assembly, 00330 Arginine and proline metabolism, 00362 Benzoate degradation, 00650 Butanoate metabolism, 01220 Degradation of aromatic compounds, 01212 Fatty acid metabolism, 00071 Fatty acid degradationl, 00380 Tryptophan metabolism, 00360 Phenylalanine metabolism. Keys for all pathways are available in **Online Resource 7**

Pathways that were underrepresented in the experimental treatments with respect to initial conditions were “Two-component system” [2020] and “ABC transporters” [2010] (sensing and active transport of DOC), “Valine, leucine and isoleucine degradation” [280], “Butanoate metabolism” [650], “Benzoate degradation” [362], “Fatty acid metabolism and fatty acid degradation” [1212 and 71] (**Figure 7**). On the other hand, pathways like “Biosynthesis of amino acids” [1230], “Ribosome” [3010], and “Purine and pyrimidine metabolism” [230 and ko240], were overrepresented in experiments with respect to initial conditions. In addition, pathways related to the metabolism of carbohydrates also increased: “Starch and sucrose metabolism” [500], “Fructose and mannose metabolism” [51], “Amino sugar and nucleotide sugar metabolism” [520]. Some of these were further overrepresented in the nutrients added condition, such as “Starch and sucrose metabolism” [500], “Other glycan degradation” [511], “Amino sugar and nucleotide sugar metabolism” [520], “Biosynthesis of amino acids” [1230], and the respiratory subsystem “Oxidative phosphorylation” [190].

## Discussion

In this work, we analyzed for the first time the exudates from *U. pinnatifida* and their effect on coastal seawater bacterial communities. Despite the drastic changes that this invasive seaweed is known to exert on invaded coastal ecosystems, until now, no information was available on the composition of *U. pinnatifida* exudates or their effect on seawater. The characteristics observed in exudates are compatible with the presence of polysaccharides and aromatic compounds, which are commonly found in exudates from other brown algae such as *Fucus vesiculosus* and *Laminaria hyperborea* (Sieburth 1969; Abdullah and Fredriksen 2004). DOM release into seawater by *U. pinnatifida* at the sampling site can be estimated based on our experimental data (1.6 ± 0.8 mg g^−1^ biomass day^−1^). Assuming conservative biomass values of ~400 g of dry weight m^−2^ (recorded in the same area in November 2006, still in early expansion phase of *U. pinnatifida* (Rechimont et al. 2013)), we can estimate a contribution of 0.64 g C m^−2^ day^−1^. This value is in the same order of magnitude of those observed for native kelps, e.g. for *Eklonia cava* in Oura Bay, Japan (0.25 to 5.8 g C m^−2^ day^−1^)(Wada et al. 2007). Integrating this value across the water column at 5 m depth, estimated daily carbon release by *U. pinnatifida* sporophytes is 0.128 mg C l^−1^ day^−1^, a significant amount with respect to the natural DOC concentration of Punta Este coastal seawater (~ 5 mg l^−1,^ as measured in this work). This is the first report of the contribution of exudates from *U. pinnatifida* to the DOC pool of coastal ecosystems invaded by this seaweed. It must be noted, however, that DOC values might be underestimated, as we observed very rapid microbial growth and optical evidence of concomitant biodegradation during exudation. On the other hand, it is possible that the experimental conditions, especially the confined containers used for the experimental systems, could affect exudation, resulting in differences with respect to natural conditions. Moreover, the quantity and quality of DOM release in *U. pinnatifida* could vary ontogenically, with temperature, and other conditions that change throughout the algae annual cycle, affecting exudation either directly and/or indirectly through changes in the algae size or metabolic state, as has been observed for the kelp *Laminaria hyperborea* (Abdullah and Fredriksen 2004) or *Eklonia cava* (Wada et al. 2007). This work was performed using mid-sized sporophytes in late winter, the period of highest biomass of the annual cycle of the kelp (Dellatorre et al. 2014). In order to estimate the contribution of *U. pinnatifida* exudates to coastal DOM and bacterioplankton throughout the year, field studies are currently being carried out, analyzing the effects of all these variables.

Exudates were composed mainly of carbohydrates, and DOM optical parameters suggested the presence of high molecular weight compounds, probably polysaccharides. Carbohydrate release was approximately 1 mg CH g^−1^ d^−1^, similar to values reported for *Fucus vesiculosus* (Sieburth 1969), but one order of magnitude higher than those observed for *Laminaria hyperborea* (0.11 mg CH g^−1^ d^−1^, (Abdullah and Fredriksen 2004). Some components of *U. pinnatifida* exudates were rapidly biodegraded, as suggested by several optical DOM proxies. In general, algal exudates are readily available C sources for free living microorganisms (Haas et al. 2011; Nelson et al. 2013), as well as for microorganisms associated to the algal surface (Bengtsson et al. 2011; Marzinelli et al. 2015) or living inside other macroorganisms such as sponges (Rix et al. 2017).

At least three fluorescent components were identified in *U. pinnatifida* exudates, two humic-like and one similar to protein or small phenolic compounds. Regarding the humic components, the C1 is ubiquitous in DOM (Stedmon and Markager 2005), whereas the C2 has been reported as marine humic-like substances (Coble 1996). Wada et al. (2008) reported that exudates of the Phaeophyceae *Ecklonia cava* were characterized by six peaks in the EEM spectra, peaks B, T, N, A, M and C. These peaks correspond to tyrosine- or protein-like (B), tryptophan- or protein-like (T), unknown (N), UV humic-like (A), visible marine humic-like (M) and visible humic-like (C) materials, respectively, following Coble and collaborators (Coble et al. 1998). This composition coincides with the characterization of *U. pinnatifida* exudates with high proportion of humic materials and the presence of the T peak. Humic-like compounds have been observed in macroalgal enriched DOM, which separated them clearly from coral-derived DOM in coral reefs, suggesting that FDOM characteristics provide accurate information for environmental assessments evaluating reef health and impact levels (Quinlan et al. 2018). Algal exudates have also been shown to be species-specific, which in turn may trigger specific bacterioplankton responses (Quinlan et al. 2018, 2019). In accordance, in our work, exudate-derived substances provoked changes in bacterial abundance, activity, and community structure in natural seawater. Exudates selected for specific bacterial populations with the corresponding metabolic capabilities. Therefore, *U. pinnatifida* exudates are a relevant source of carbon and its associated microorganisms, with the potential to trigger profound changes in the coastal ecosystem in this area.

The capabilities for processing polysaccharides vary widely among microbial populations. Multiple factors such as substrate quantity and quality, accessibility, and microbial capabilities including sensing the substrates and producing the correct enzymes under specific environmental conditions, are part of this complex process, giving rise to differences that can be observed at the community level (Arnosti 2014). Although the effect of macroalgae exudates on coral reefs communities has been studied in detail (see Quinlan et al. 2019, and references therein), limited information is available on the effect of this abundant source of carbon on bacterioplankton. The bacterial populations stimulated by the exposure to algal exudates in this work are probably chemoorganotrophs, specialized in the metabolism of algal carbohydrates. For example, the Flavobacteria are abundant members of marine microbial communities, with either planktonic or algal-associated lifestyles, and known to be key players in the biodegradation of micro and macroalgal-derived complex organic matter through a battery of specialized extracellular enzymes (Bauer et al. 2006; Alonso et al. 2007; Fernández-Gómez et al. 2013; Mann et al. 2013; Chernysheva et al. 2019).

Another group found to be enriched were members of the *Kiritimatiellaeota* (previously *Verrucomicrobia* subdivision 5). This is an emergent phylum, its members only just beginning to be studied. The genera known so far are anaerobic, saccharolytic organisms which obtain energy from fermentation. The first cultured representatives of this group, isolated from a hypersaline lake and from marine sediments, also stand out for their capability of degrading complex sulfated polysaccharides such as fucoidans, as well as glycoproteins (Spring et al. 2016; Sackett et al. 2019; van Vliet et al. 2019). Interestingly, the role of members of this new phylum as well as other related PVC groups could be important in hypoxic environments such as bottom waters and surface sediments, where organic matter accumulates (Cardman et al. 2014). Another group of microorganisms that were observed in exudate-enriched systems were facultatively anaerobes typical of aquatic environments, like the genus *Spirochaeta* (Spirochaetes) and *Propionigenium* (Fusobacteria). These organisms are capable of performing fermentative metabolism of small carbohydrates (Leschine et al. 2006; Gupta and Sethi 2014) and, particularly the Fusobacteria, are commonly found in suboxic environments such as sediments and in mucous membranes and guts of animals, including algal-feeding marine invertebrates (Janssen and Liesack 1995; Hakim et al. 2016; Yao et al. 2019). Members of the genus *Arcobacter,* which also shifted in abundance and increased with nutrients addition, are characterized by an unusually wide range of habitats, although mainly heterotrophic and host-associated (Waite et al. 2017; Fanelli et al. 2019). It must be noted that the majority of these groups might have not been represented in plate counts, which favor aerobic microorganisms.

Overall, the results of this study show evidence of a highly integrated and coordinated microbial community in seawater exposed to exudates from *U. pinnatifida*, where some members likely specialize in degradation of complex polysaccharides, and are accompanied by fermentative bacteria probably able to proliferate on their degradation products.

Some of the bacterial populations found to be favored by exudates, such as *Arcobacter* and *Vibrio*, are known for their pathogenic properties, raising concerns about potential environmental and ecotoxicological consequences of the invasion by *U. pinnatifida*. For example, members of the genus *Arcobacter* have been classified as emergent human pathogens (Snelling et al. 2006; Fanelli et al. 2019). They have been detected in shellfish for human consumption (Collado et al. 2009; Laishram et al. 2016). Moreover, strains isolated from shellfish, which are considered their marine reservoir, have been shown to display both antibiotic resistance and virulence factors (Fanelli et al. 2019). On the other hand, *Vibrio* populations are common pathogens of bivalves, affecting both wild and cultured populations (Beaz Hidalgo et al. 2010; Dubert et al. 2017). Notably, the abundances of ASVs assigned to *Arcobacter* and *Vibrio* further increased in the nutrient-added condition, highlighting the role of nutrient availability in the development of potentially pathogenic bacteria. Nutrients have been shown to increase the severity of coral disease, via differential utilization of nutrients by pathogenic *Vibrio* populations, which in turn increased their fitness and virulence (Bruno et al. 2003). Nutrients also enhance exudation in coralline algae (Quinlan et al. 2018), a condition that, together with microbial communities stimulation, could set a positive feedback loop. This scenario may be more significant in a low energy environment such as the coasts of Nuevo Gulf.

The metabolic potential prediction performed from the bacterial community structure data reflected a shift from the typical bacterioplankton assemblages dominated by generalists able to sense, incorporate and utilize small compounds (amino acids, glucose and fatty acids) (Mou et al. 2008; Poretsky et al. 2010), towards other microbial groups with higher metabolic rates, adapted to higher concentrations and/or to the presence of high molecular weight carbon compounds, such as polymers of sugar and other glycans. The higher number of ribosomal genes predicted in the latter group of organisms reflects differences in copy numbers that have also been linked to a copiotrophic life strategy in bacteria (Klappenbach et al. 2000). Therefore, our results suggest that the shifts observed in these communities were indeed associated to the presence of exudates, which favored the emergence of populations with different ecophysiological niches (Cottrell and Kirchman 2000; Alonso and Pernthaler 2006).

## Conclusion

The impact of invasive kelp species can be analyzed at multiple levels, among which physicochemical and microbial aspects are of paramount importance, given the profound effects they can have on ecosystem functioning. The results of this work highlight the mechanisms by which two human-associated stressors, DOM released by an invasive algal species and nutrient pollution, can affect seawater physicochemical properties and microbial communities, modifying the natural bottom up control in coastal food webs with unforeseen consequences on the coastal ecosystem and its services. Further work is currently being carried out in this direction, addressing changes in various chemical and microbiological aspects in coastal environments of Patagonia affected by *U. pinnatifida* throughout its yearly cycle.

## Supporting information

Online Resource 1

Online Resource 2

Online Resource 3

Online Resource 4

Online Resource 5

Online Resource 6

Online Resource 7

## Declarations

### Funding (information that explains whether and by whom the research was supported)

All authors are staff members of The Argentinean National Research Council (CONICET). This work was supported by CONICET (Grant PIP2014-2016 No 112-200801-01736) and ANPCyT (Grant PICT-2015-2102, PICT-2016-2101 and PICT-2018-0903).

### Ethics approval

(include appropriate approvals or waivers): not applicable

### Consent to participate

(include appropriate statements): not applicable

### Consent for publication

(include appropriate statements): all authors have read and approved the final version of this manuscript. This work has not been sent to any other journal for publication.

### Availability of data and material

(data transparency): the sequence information has been deposited in NCBI Short Sequence Archive (SRA) under BioProject ID: PRJNA622742

### Code availability

(software application or custom code): the scripts used for processing sequences have been included as Online Resource.

## Acknowledgements

All authors are staff members of The Argentinean National Research Council (CONICET). This work was supported by CONICET (Grant PIP2014-2016 No 112-200801-01736) and ANPCyT (Grants PICT-2016-2101 and PICT-2018-0903).

