## Supplementary material for "*Undaria pinnatifida* exudates trigger shifts in seawater chemistry and microbial communities from Atlantic Patagonian coasts": Online Resource 1

**Supplementary methods**

The analysis of chromophoric dissolved organic matter (CDOM) and its fluorescent fraction (FDOM) was assessed through absorption and fluorescence spectroscopy, respectively. Absorption spectra of filtered (0.22 µm) water samples were obtained from 200 to 800 nm at 1 nm intervals with a UV/Vis spectrophotometer (Shimadzu, UV1800), using a quartz cuvette (10 cm path length). Absorbance data was converted to absorption coefficients (*a_λ_*) following Helms et al. (2008). The spectral slope for the interval 275-295 nm (S_275-295_) was calculated and applied as a proxy of molecular weight/size and degree of aromaticity with higher S values indicating low molecular weight material and/or decreasing aromaticity (Helms et al., 2008; Hansen et al., 2016).

Excitation-emission matrices (EEMs) of FDOM were obtained by scanning the filtered water samples in a spectrofluorometer Perkin Elmer LS55 equipped with a 150-W Xenon arc lamp and a Peltier temperature. The EEMs were collected at excitation wavelengths, from 240 to 450 nm, every 5 nm, with a slit width of 15 nm. Emission wavelengths were collected from 300 to 598.5 nm every 0.5 nm, with a slit width of 15 nm at a scan speed of 1500 nm min^-1^ (integration time of 0.2 sec). The EEMs were corrected for the inner filter effect and normalized to the area under the Raman peak of the blank (MilliQ water) at 350 nm using the FDOMcorr toolbox (Murphy et al, 2010). The resulting data was expressed in Raman units.

Three fluorescent indexes were calculated to characterize the FDOM fraction following Hansen et al (2016). The humification index (HIX) was computed as the sum of emission intensities between 435 and 480 nm divided by the sum of emission intensities between 300 and 345 nm at 254 nm excitation and used as a proxy of humic content. The biological index (BIX) was calculated as the ratio of emission intensity at 380 nm divided by 430 nm at excitation 310 nm and applied as an indicator of autotrophic productivity. BIX values >1 are indicative of recently produced DOM of autochthonous origin. The fluorescence index (FI) was calculated as the emission ratio of 470 and 520 nm at excitation 370 nm and applied to describe the relative contribution of terrestrial and microbial sources .

**Parallel factor analysis on FDOM data**

Parallel factor analysis (PARAFAC) was performed using a set of 46 EEMs collected from both experiments. The Raman and Rayleigh scatter were removed from the analysis according to Stedmon and Bro (2008). An exploratory analysis using non-negative constraints was performed in order to identify outliers (leverage < 0.5). The components were validated using split-half and the best model fit was obtained using 10 random initializations (Stedmon and Bro, 2008; Murphy et al., 2013). The DOMFLUOR and DrEEM toolboxes developed for MATLAB (MATLAB R 2014a) were used to process the EEMs and to perform the PARAFAC (Stedmon and Bro, 2008; Murphy et al., 2010; 2013). The components validated in the study were contrasted (similarity score >0.98) with the components listed in the OpenFluor database (Murphy et al., 2014).

Table caption: Characterization of the FDOM components modelled through PARAFAC. Component´s matches presented are exclusively from marine DOM.

| **Component number (Ex/Em)** | **Component EEM** | **Component loading** | **Match reference (Openfluor)** |
| --- | --- | --- | --- |
| C1: 320(240)/394 | 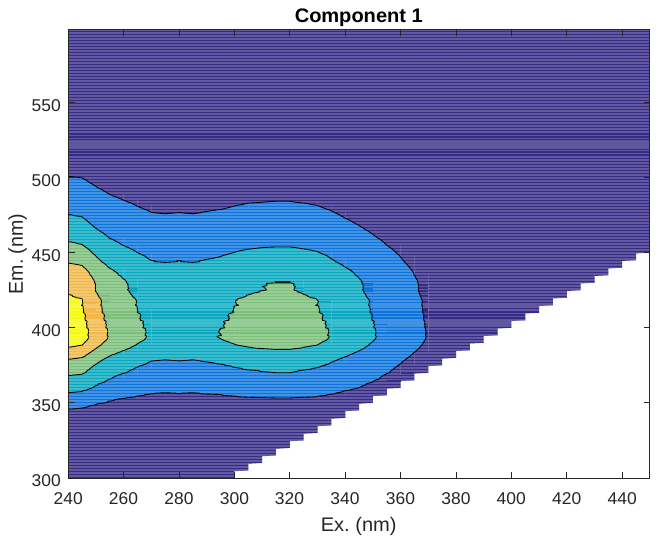 | 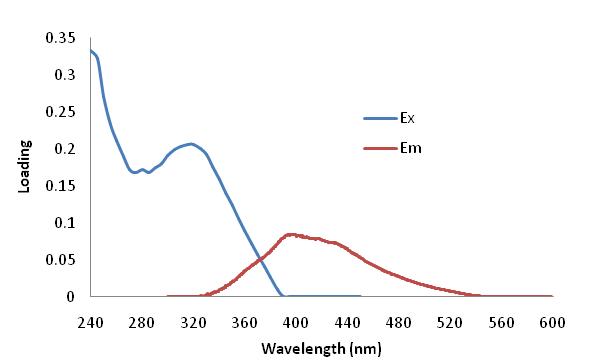 | Catalá et al., 2015 (C2)  Cawley et al., 2012(C1)  Wünsch et al., 2018 (C1) |
| C2: 375(250)/350 | 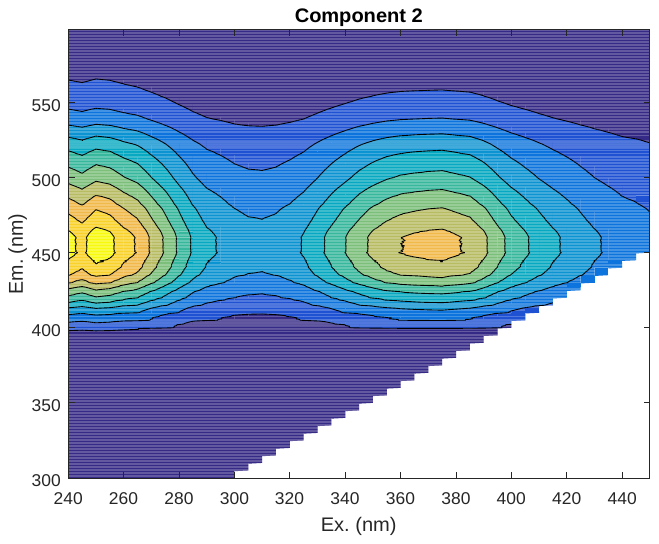 | 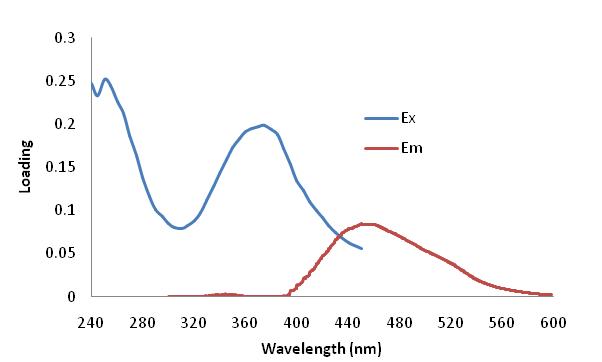 | Yamashita et al., 2010 (C1)  Derrien et al., 2019 (C2) |
| C3: 280/341.5 | 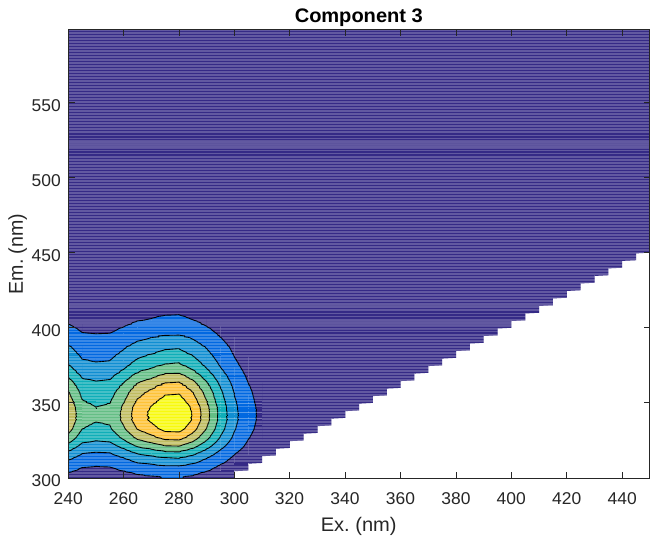 | 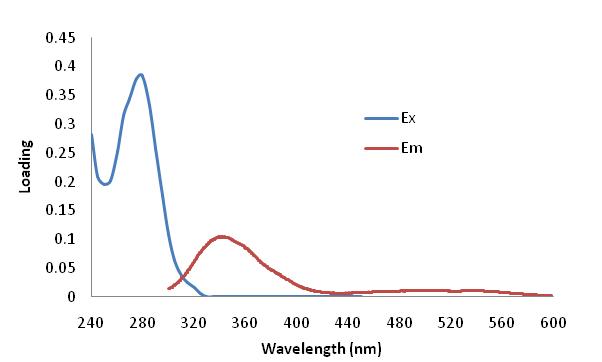 | Osburn et al., 2012 (C5)  Osburn et al 2016(C3) |

Catalá, T. S. et al., 2015. Turnover time of fluorescent dissolved organic matter in the dark global ocean. Nat. Commun. 6: 5986.

Cawley et al., 2012. Characterising the sources and fate of dissolved organic matter in Shark Bay, Australia: a preliminary study using optical properties and stable carbon isotopes, Marine and Freshwater Research 63: 1098-1107.

Derrien, M., Retelletti Brogia, S. Goncalvez-Araujo, R. 2019. Characterization of aquatic organic matter: Assessment, perspectives and research priorities. Water Research 163: 114908.

Wünsch, U. J., Geuer, J. K., Lechtenfeld, O. J., Koch, B. P., Murphy, K. R., & Stedmon, C. A. 2018. Quantifying the impact of solid-phase extraction on chromophoric dissolved organic matter composition. Marine Chemistry 207, 33-41.

Yamashita, Y., Cory, R.M., Nishioka, J., Kuma, K., Tanoue, E., Jaffe, R., 2010. Fluorescence characteristics of dissolved organic matter in the deep waters of the Okhotsk Sea and the northwestern North Pacific Ocean. Deep-Sea Res II 57, 1478-1485.

Osburn, C.L., Handsel, L.T., Mikan, M.P., Paerl, H.W., and Montgomery, M.T., 2012. Fluorescence Tracking of Dissolved and Particulate Organic Matter in a river-dominated estuary. Environ Sci Technol. 46(16):8628-8636.

Osburn, C. L., Handsel, L. T., Peierls, B. L., & Paerl, H. W., 2016. Predicting sources of dissolved organic nitrogen to an estuary from an agro-urban coastal watershed. *Environmental science & technology*, *50*(16), 8473-8484.
