## Supplementary material for "*Undaria pinnatifida* exudates trigger shifts in seawater chemistry and microbial communities from Atlantic Patagonian coasts": Online Resource 3

**Figure caption.** Multiple correlation analysis of data from experiment 1 (incubation systems with *U. pinnatifida* in seawater). BIX: biological index. S275-295: spectral slope calculated for the decrease in absorbance between 275 and 295 nm. FI: fluorescence index. HIX: humification index. FDOM components C3(T), C1(A+M), C2(A+C):. DOC: dissolved organic carbon concentration. CH: Total carbohydrates concentration. A_440_: Absorbance at 440 nm.

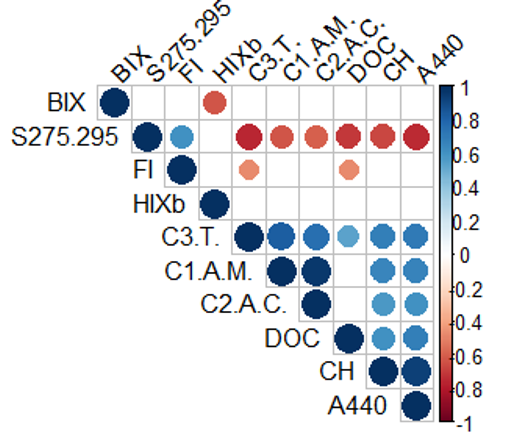

**Table caption.** Correlation between response variables measured in Exp 1 and the three principal components obtained in the PCA.

|  | PC1 | PC2 | PC3 |
| --- | --- | --- | --- |
| CH | **0.866** | -0.235 | 0.211 |
| DOC | **0.722** | -0.187 | -0.374 |
| S275.295 | **-0.878** | -0.029 | 0.307 |
| HIX | 0.232 | **-0.875** | 0.282 |
| BIX | -0.161 | **0.824** | 0.141 |
| a440 | **0.911** | -0.202 | 0.073 |
| FI | -0.506 | -0.084 | **0.733** |
| C1 (A+M) | **0.833** | 0.284 | 0.399 |
| C2 (A+C) | **0.792** | 0.233 | 0.376 |
| C3 (T) | **0.872** | 0.424 | -0.014 |
| Eigenvalue | 5.290 | 1.897 | 1.220 |
| Variance | 52.897 | 18.973 | 12.203 |
| Cumulative Variance | **52.897** | **71.870** | **84.073** |
