## Supplementary material for "*Undaria pinnatifida* exudates trigger shifts in seawater chemistry and microbial communities from Atlantic Patagonian coasts": Online Resource 4

**Table caption.** Nutrient content in experimental systems of seawater incubations with exudates from *U. pinnatifida* (experiment 2). Concentrations are expressed in micromoles per liter (μM) and expressed as mean ± 1 standard deviation. EX: exudate, EXN: exudate plus nutrients, INC: incubation control. Exp_time: experiment (*2*: exposure of exudate to fresh seawater) followed by day of experiment (1 to 5). sd: standard deviation. n: number of samples per treatment. na: not applicable (n=1).

| **Treatment** | **Time**  **(days)** | **n** | **NH_4_**  **(μM)** | **NH_4_-sd** | **NO_3_NO_2_**  **(μM)** | **NO_3_NOsd** | **PO_4_**  **(μM)** | **PO_4_-sd** | **SiO_3_**  **(μM)** | **SiO_3_-sd** |
| --- | --- | --- | --- | --- | --- | --- | --- | --- | --- | --- |
| EX | 0 | 3 | 7.8 | 3.3 | 7.6 | 1.6 | 1.4 | 0.1 | 6.2 | 0.6 |
| EX | 1 | 3 | 8.0 | 4.9 | 7.3 | 2.3 | 1.7 | 0.0 | 6.4 | 0.5 |
| EX | 3 | 3 | 12.2 | 5.2 | 6.4 | 2.7 | 2.1 | 0.1 | 5.6 | 0.4 |
| EX | 5 | 3 | 11.3 | 7.9 | 7.6 | 2.9 | 2.1 | 0.1 | 5.7 | 0.4 |
| EXN | 0 | 3 | 127.8 | 7.1 | 6.8 | 2.7 | 12.4 | 0.1 | 6.7 | 0.4 |
| EXN | 1 | 3 | 131.1 | 7.5 | 13.3 | 5.8 | 12.8 | 0.6 | 6.1 | 0.1 |
| EXN | 3 | 3 | 161.8 | 37.1 | 4.8 | 2.0 | 12.7 | 0.1 | 6.6 | 0.7 |
| EXN | 5 | 3 | 139.2 | 3.8 | 6.3 | 2.3 | 12.6 | 0.3 | 11.1 | 8.7 |
| INC | 0 | 1 | 3.8 | na | 13.7 | na | 1.7 | na | 6.2 | na |
| INC | 1 | 1 | 1.9 | na | 12.5 | na | 1.6 | na | 5.7 | na |
| INC | 3 | 1 | 1.9 | na | 5.8 | na | 1.8 | na | 6.6 | na |
| INC | 5 | 1 | 1.9 | na | 5.8 | na | 1.9 | na | 5.8 | na |

**Table caption.** Characteristics of experimental systems containing fresh seawater enriched with *U. pinnatifida* exudate used for microbial community structure analysis by molecular methods. DOC: dissolved organic carbon concentration; DO: dissolved oxygen concentration; CH: carbohydrates; CFU: colony forming units. Values are expressed as mean ± 1 standard deviation. n= number of biological replicates. nd: not determined

| **Treatment** | **Time**  **(days)** | **n** | **DOC**  **(mg/L)** | **CH**  **(mg/L)** | **pH** | **DO**  **(%)** | **Bacterial abundance (CFU/ml)** |
| --- | --- | --- | --- | --- | --- | --- | --- |
| EX | 0 | 1 | 42 | 14.9 | 7.51 | 50 | nd |
| EX | 5 | 3 | nd |  |  | 35 ± 10 | 5.3±0.08 x10 ^5^ |
| EX | 7 | 3 | 61 ± 2 | 13.4 ± 0.3 | 6.48 ± 0.07 | 14 ± 3 | 3.0±0.12 x10 ^5^ |
| EXN | 5 | 3 | nd |  |  | 29 ± 1 | 3.00±0.9 x10 ^6^ |
| EXN | 7 | 3 | 41 ± 3 | 12.8 ± 0.6 | 6.37 ± 0.10 | 16 ± 1 | 2.5±0.1 x10 ^6^ |
