## Supplementary material for "*Undaria pinnatifida* exudates trigger shifts in seawater chemistry and microbial communities from Atlantic Patagonian coasts": Online Resource 5

Experiment 2

Correlation matrix

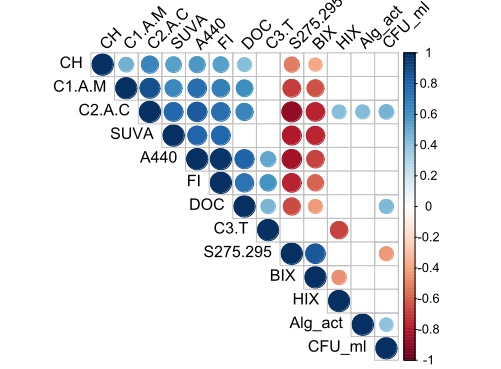

Table PCA Experiment 2

| **Variable** | **PC1** | **PC2** | **PC3** |
| --- | --- | --- | --- |
| Alg_act | 0.40 | -0.23 | -0.28 |
| CFU/ml | 0.43 | -0.45 | **0.59** |
| CH | **0.65** | -0.18 | 0.14 |
| DOC | **0.73** | 0.19 | 0.56 |
| S275.295 | **-0.93** | 0.10 | 0.07 |
| HIX | 0.31 | **-0.84** | 0.01 |
| BIX | **-0.80** | 0.20 | 0.37 |
| A440 | **0.94** | 0.32 | 0.05 |
| SUVA | **0.83** | 0.10 | -0.44 |
| FI | **0.88** | 0.43 | 0.01 |
| C1(A+M) | **0.84** | 0.00 | 0.02 |
| C2 (A+C) | **0.96** | -0.17 | -0.01 |
| C3 (T) | 0.19 | **0.94** | 0.06 |
| Eigenvalue | 6.9 | 2.29 | 1.11 |
| Variance | 53.08 | 17.62 | 8.54 |
| CumulativeVariance | 53.08 | 70.7 | 79.24 |
